# Differential gene expression in brain and liver tissue of Wistar rats after rapid eye movement sleep deprivation

**DOI:** 10.1101/515379

**Authors:** Atul Pandey, Ryan Oliver, Santosh K Kar

## Abstract

Sleep is essential for the survival of most living beings. Numerous researchers have identified a series of genes that are thought to regulate “sleep-state” or the “deprived state”. As sleep has significant effect on physiology, we believe that lack of sleep or particularly REM sleep for a prolonged period would have a profound impact on various body tissues. Therefore, using microarray method, we sought to determine which genes and processes are affected in the brain and liver of rats following 9 days of REM sleep deprivation. Our findings showed that REM sleep deprivation affected a total of 652 genes in the brain and 426 genes in the liver. Only 23 genes were affected commonly, 10 oppositely and 13 similarly across brain and liver tissue. Our results suggest that 9-day REM sleep deprivation differentially affects genes and processes in the brain and liver of rats.

**Highlights of the study:**

➢ Gene expression profile of brain and liver tissues of rats was analyzed using microarray technique following 9 days of REM Sleep deprivation.
➢ Many of the genes involved in essential physiological processes, such as protein synthesis and neuronal metabolism are affected differently in the brain and liver tissue of rats after 9-day REM sleep deprivation.

## 1. Introduction

Sleep is a universal phenomenon but still we lack fundamental knowledge of its overall functions and purpose. However, most comparative sleep data exist for terrestrial vertebrates, with much less known about sleep in invertebrates [1]. Though, recently the scientific community has sought to characteristic sleep in non-mammalian species like the fruit fly (*Drosophila melanogaster*)[2–4], the zebrafish (*Danio rerio*) [5–7], the nematode (*Caenorhabditis elegans*) [8], and bees (*Apis mellifera*, and *Bombus terrestris*) [9–12]. Prolonged sleep deprivation is fatal in many animals studied so far except pigeons and several studies have sought to address how sleep promotes survival in rodents and primates [13–16]. Despite the lack of general knowledge regarding the functions of sleep, loss of sleep has been shown to drastically alter physiology of many animals studied thus far [17–19]. The degree of physiological changes that sleep loss brings about and the fatality often varies depending upon the nature and duration of sleep deprivation [20,21]. Many theories have been proposed to explain the evolutionary significance and functions of sleep, which includes “null” and “synaptic plasticity” theories [22,23]. Recent advancements in sleep research has shed light on two major functions of sleep-reducing synaptic potentiation and waste clearance mediated by glymphatic system [24–26]. Thus, sleep seems to have specific, overarching functions for all species that depend on it [19]. While a single characterization cannot be ascribed to sleep, numerous studies links its loss to detrimental effects on metabolism, behavior, immunity, cellular functions and hormonal regulations across species [27–30]. Thus, we may suggest that sleep is generally necessary, and most living beings cannot be deprived of it for a long time. There are some mechanisms that are associated with behavioral plasticity which are dependent on sociality or physiological state in regard to sleep regulation [31,32]. Also, in *Drosophila*, not all stages of sleep are necessary for basic survival but questions relating to the critical functions of sleep, plasticity, and its overall importance are still being explored [31].

REM sleep is an essential part of sleep and is present only in avians and mammals, with the exception of reptiles, in which REM sleep has only been recently discovered [33]. Unearthed thus far, the functional aspects of REM sleep includes mainly memory consolidation, brain maturation, muscle re-aeration, special memory acquisition, and maintenance of general physiological mechanisms of the body [34–40]. In the brain, REM sleep is involved in the reorganization of hippocampal excitability, pruning and maintenance of new synapses during development, and learning & memory consolidation [41–43]. Some recent studies also suggest that lack of REM sleep may cause cell death of somatic cells and neurons [44–46]. Outside of the brain, deprivation of the REM sleep was found to be associated with acute phase response in the liver, increased synthesis of pro-inflammatory cytokines such as IL1β, IL-6, and IL-12, and an increase in liver enzymes, alanine transaminase and aspartic transaminase [47]. In addition, REM sleep deprivation induces the production of reactive oxygen species (ROS), caused inflammation [48] and increase in nitric oxide (NO) in hepatocytes, along with an increase in sensitivity to oxidative stress by the hepatocytes [49]. REM loss also affected the weight and content of nucleic acid in liver [50]. REM loss was also found further associated with oxidative stress and liver circadian clock gene expression [51]. Elevated increase in metabolic rate and UCP1 gene expression is reported in response to chronic REM sleep loss in brown adipose tissue of rats [52]. Recently, REM sleep loss is found associated with blood-brain barrier function regulation and metabolic changes [53,54].

On genomic level, the reduction of gene expression related to energy metabolism (e.g. glucose type I transporter Glut1), growth (e.g. Bdnf), vesicle fusion and many other metabolic processes has been found affected by sleep [55,56]. Another study detected a decrease in GluR1-containing AMPA receptor (AMPAR) levels during sleep, as well as a decrease in AMPAR, CamKII and GSK3beta phosphorylation [57]. The Synaptic Homeostasis Hypothesis (SHY), which postulates that wakefulness and sleep are linked to a net increase and decrease in synaptic strength, is supported by these findings [58,59]. In rodents, synaptic plasticity-related expression levels of immediate early genes (IEG), such as *Egr1, Arc* and *Fos*, were found to decrease from wakefulness to sleep [56,60–68]. Some of the genes related to synaptic plasticity theory e.g., *Arc, Bdnf, Creb1, Egr1, Fos, Nr4a1, Camk4, Ppp2ca*, and *Ppp2r2d* were studied in detail for short wave sleep and REM sleep [69]. It has been proposed that some of these genes, such as *Arc* and *Egr1*, play a key role in long term potentiation (LTP) [60,61,63–65,67,70–73]. It is suspected that other genes are also important for LTD, such as *Ppp2ca* and *Ppp2r2d*, which code for subunits of *PP2A* [74]. While, REM sleep deprivation in the rat dorsal hippocampus has been shown to decrease LTP, synaptic transmission, protein levels of the glutamate receptor, and activation of ERK / MAPK [75].

In the present study, we compared the effect of prolonged REM sleep loss in the brain and liver in order to compare and contrast the effects that occur simultaneously on these vital organs, as previous studies had indicated that REM sleep may have drastic effects on the liver [45,47,49]. To address this, we used a microarray technique to compare gene expression and identify the processes affected in the brain and liver of a given subject after REM sleep deprivation for 9 days. Microarray is a valuable tool for measuring the dynamics of gene expression in a biological system and can be used to measure the differences in gene expression profile of different tissues under the same physiological conditions [19,76,77]. Most sleep studies that involved microarray analysis have been performed in brain, although recently, research has indicated that other organs may also play a crucial role [55,78–83]. We first hypothesized that prolonged REM sleep loss will differentially affect genes and associated processes in the brain and liver. Secondly, we hypothesized that REM sleep loss would affect functions related to synaptic potentiation and maintenance in the brain, and metabolism and immune response to infection related mechanisms, in the liver.

Previous studies involving analysis of microarray returned many genes which were associated with the GO term, potentiation of synaptic plasticity, which largely supports the ‘synaptic homeostasis theory’ [55,84]. In the cerebral cortex of the mouse and, to a lesser degree, hypothalamus, genes encoding proteins of various biosynthetic pathways for heme, protein, and lipid are upregulated throughout sleep [81]. Throughout sleep, a significant number of genes encoding the structural constituents of the ribosomes, translation-regulation activity, and formation of transfer RNA (tRNA) and ribosome biogenesis are also upregulated. Genes whose expression gradually increases during sleep include those that encode for several cholesterol-synthesis pathway enzymes, proteins involved in the uptake of cholesterol, the transport of transcription factors, and chaperones that regulated the transcription of genes associated with cholesterol [81]. Prolonged wakefulness results in the fruit fly results in the downregulation of several genes involved in protein production [83]. Sleep deprivation in mice causes a decrease in the expression of genes in the cerebral cortex and hypothalamus, which encode proteins that are involved in key pathways of carbohydrate metabolism, energy production, tricarboxylic acid (TCA) anabolism, and various metabolic pathways (lipid, aldehyde, amine synthesis) [81]. Further, microarrays have shown that there are transcript level variations in many genes involved in the regulation of reactive oxygen species (ROS), including heme oxygenase, superoxide dismutase, and catalase, in patients with obstructive sleep apnea [85]. The dopamine receptor-signaling pathway regulating sleep, learning, and its plasticity are well known [86,87]. Sleep disorders and sleep deprivation have been correlated with dopaminergic, cholinergic, and GABAergic regulation of synaptic transmission, each of which were terms that were significantly enriched for genes that were downregulated in our study [56,88–91]. A recent microarray analysis involving mice shows that *Hspa5* gene expression increases not only in the brain but also in the liver as sleep deprivation increases [82]. Overall, currently, however, there is little knowledge available about how sleep including REM, its loss and the prolonged wakefulness affects expression of genes in peripheral tissues, an area that is open for future research. Our current study fits nicely here to answer many REM sleep loss related questions comparing microarray dataset between brain and liver and provide unique dataset for future research.

## 2. Material and Methods

Male Wistar rats, weighing between 220-260 grams, were used for this study. Animals were housed with 12:12hrs L: D cycle (7:00 am lights on) and provided with food and water *ad libitum*. All experiments were carried out in compliance with the Institutional Animal Ethics Committee of the University.

### 2.1. REM sleep deprivation procedure

Rats were REM sleep-deprived for nine consecutive days by using the flower pot method [92,93]. Subjects were kept on a relatively small, raised platform (6.5 cm in diameter) and surrounded by water. While, for the sham control (large platform control, LPC) animals are kept on a larger platform (12.5 cm in diameter) under similar conditions of experimental group. REM sleep-deprived animals could sit, crouch, and have a NREM-related sleep on this platform. However, due to muscle atonia during REM sleep, they are unable to have REM sleep on the small platform. Upon entering REM sleep, subjects fell into water in order to disrupt the entirety of its cycle. Throughout our previous studies, there were no differences between the cage control (animals kept in cages) and LPC control group of rats, and thus only the LPC control group is referred to as the “**control**” group [47,49]. Rats were sacrificed between 10 a.m. and 12 p.m. on day 9 and the total brain and liver were harvested and flash-frozen in liquid nitrogen for further analysis.

### 2.2. RNA extraction and quality analysis

Total RNA was isolated from the entire brain and liver samples using standard protocol. Rats were anesthetized with isoflurane, and brain and liver samples were immediately dissected and frozen in liquid nitrogen. We isolated total RNA from the whole brain and liver of each animal using Trizol methods (Gibco-BRL, Gaithersburg, MD, USA), as directed by the manufacturer. The concentration of total RNA was measured using Nanodrop and quality analyzed using Bioanalyzer.

### 2.3. Microarray: labeling, hybridization, and data analysis

An equal amount of total RNA from the brain and liver were collected and sent to Ocimum Biosolutions (USA) genomics facility for microarray analysis. Affymetrix Rat Gene 1.0 ST Arrays containing more than 7000 annotated sequences and 18,000 expressed sequence tags (ESTs) were used. The Affymetrix Gene Chip Expression Technical Manual (Affymetrix Inc., Santa Clara, CA, USA) was used for marking, hybridization, and expression analysis of microarrays, according to previous methods [79]. The data analysis was performed using Affymetrix Expression Console and Programming Language-R [94,95].

### 2.4. Gene Ontology analysis

Functional annotations of differentially expressed genes were obtained from the Gene Ontology Consortium database, based on their respective biological process, molecular function and cellular component [96]. Overrepresentation analysis, using a single-tailed Fisher exact probability test, based on the hypergeometric distribution, was used and significant GO terms were stored (*p*< 0.05).

### 2.5. Pathway analysis

Pathway analysis of microarray data was performed using the Kyoto Encyclopedia of Genes and Genomes (KEGG) software. Several biochemical pathways are identified by physiological processes documented in the KEGG databank. Since rat species-specific functional gene annotations are still few for several biological processes, general pathways, pathways of other organisms, and species-specific pathways were combined for a comprehensive analysis. We used the KEGG map pathway to visualize the maximal impact of REM sleep loss on highly up-regulated genes involved in protein translation processes.

### 2.6. Validation of array expression with Real-Time quantitative qPCR

Following analysis of microarray, a group of genes were selected for validation by qPCR, based on their degree of change in expression. We tested for correlation between the effects of the microarray and qPCR, and statistical significance was calculated (**Fig. 1**). The microarray data used for the correlation was input as Log2 ratio of the weighted average of each gene per composite array for all subjects. For qPCR, we used the mean Log2 ratio value stated by qPCR of each subject. Six transcripts were selected for validation of microarray analysis using RT-PCR (**Supplementary Table, ST-1**). Controls were used to rule out the effect of any confounding variables. We tested the respective mRNA levels with RT-PCR. Samples obtained from liver and brain tissue were frozen and stored separately at −80°C before mRNA was quantified. Total RNA was isolated using Trizol methods and re-transcribed using the ABI reverse transcription kit (Applied Biosystems, Catalog number: 4368814). TaqMan gene expression Master Mix (Applied Biosystems, Catalog Number: 4369016) and probes (Applied Biosystems, **Supplementary Table, ST-1**) used for quantitative analysis of mRNA. Each cDNA sample was analyzed in triplicate. The RT-PCR reactions for all focal genes and Glyceraldehyde 3-phosphate dehydrogenase (GAPDH) were measured from the same cDNA sample and loaded onto the same 96-well analysis plate. We quantified the gene levels using 2 ^−^ΔΔCt methods and GAPDH was used as a reference, control gene for expression level normalization. Expression validation experiments were performed on the basis of five rats per group.

**Figure 1.**
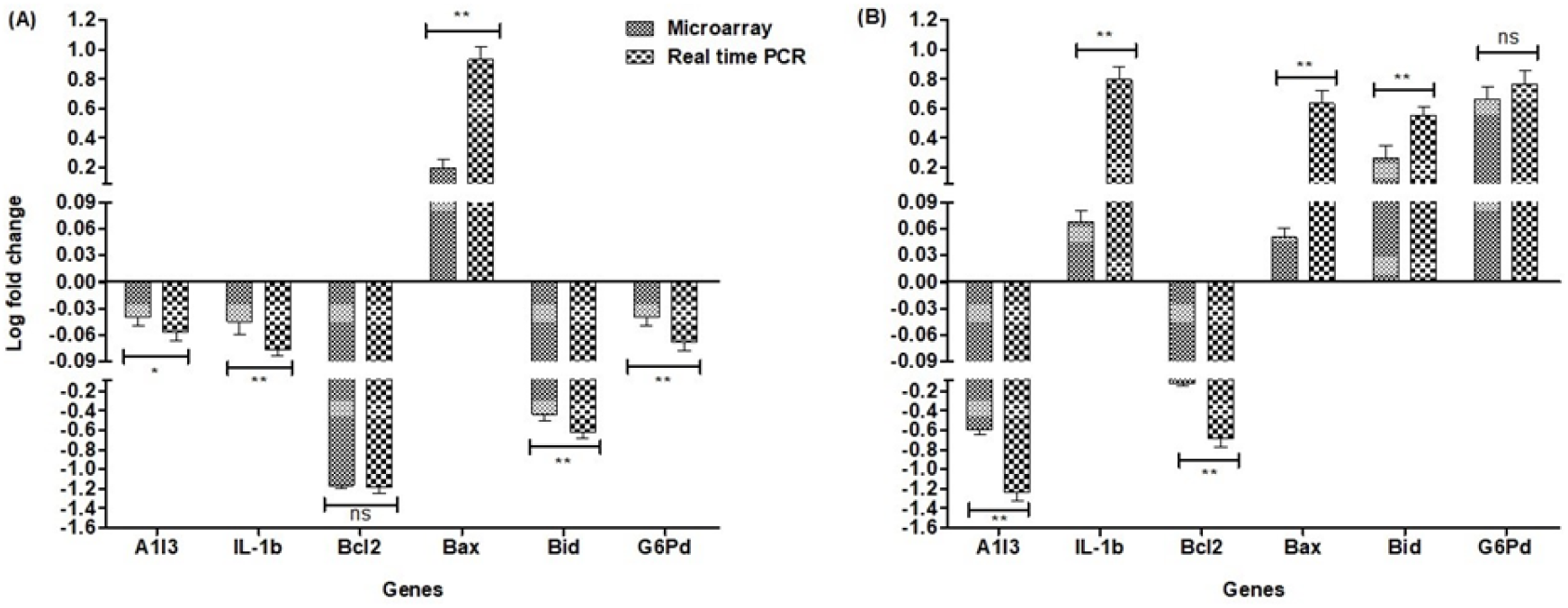
Relative expression of six candidate genes from the brain and liver tissue using real-time PCR and Microarray. (A) the comparative expression of genes in the brain; (B) the comparative expression of genes in the liver. Relative gene expressions were normalized by comparison with the expression of the Glyceraldehyde 3-phosphate dehydrogenase (GAPDH) gene, while results were analyzed using the 2−ΔΔCT method. For each gene, all RT-qPCRs used five biological replicates, with three technological replicates per experiment. The non-parametric Mann-Whitney U test was used to compare the pairwise expression of the microarray and the RT-PCR expression for the respective genes. We evaluated the normality of the data using the Kolmogorov-Smirnov normality test. Error bars indicate a ±SE value.

### 2.7. Statistics

The results of qRT-PCR are presented as a mean of ± SE. We used the Kolmogorov – Smirnov normality test to estimate the normality of the data. The Mann-Whitney U test was used to compare the pairwise expression of the microarray and RT-PCR expression for the respective genes used for liver and brain validation. The array experiments were analyzed, maintaining a *p*<0.5 significance level. The KEGG bioinformatics map and diagrams were built based on an analysis of semantic similarity of terms, using Wang’s method. Visualization of connectivity in network plots were designed in R using cluster Profiler package [97,98]. All statistical analyzes considered *p*< 0.05 to be significant and were performed and plotted using Sigma 8.0 and 12.0, Graph Pad 5.1 and Origin 6.0 software.

## 3. Results

### 3.1. General results

In the current analysis, we used the Affymetrix Rat Gene 1.0 ST Array and data analysis was performed using Affymetrix Expression Console and R-software. A total of 311 up-regulated genes (**Supplementary Table, ST-2A**) and 341 down-regulated genes (**Supplementary Table, ST-2B**) have been found in the brain. In contrast, 209 up-regulated genes (**Supplementary Table, ST-2C**) and 217 genes were down-regulated (**Supplementary Table ST-2D**) in the liver (**Fig. 2**).

**Figure 2.**
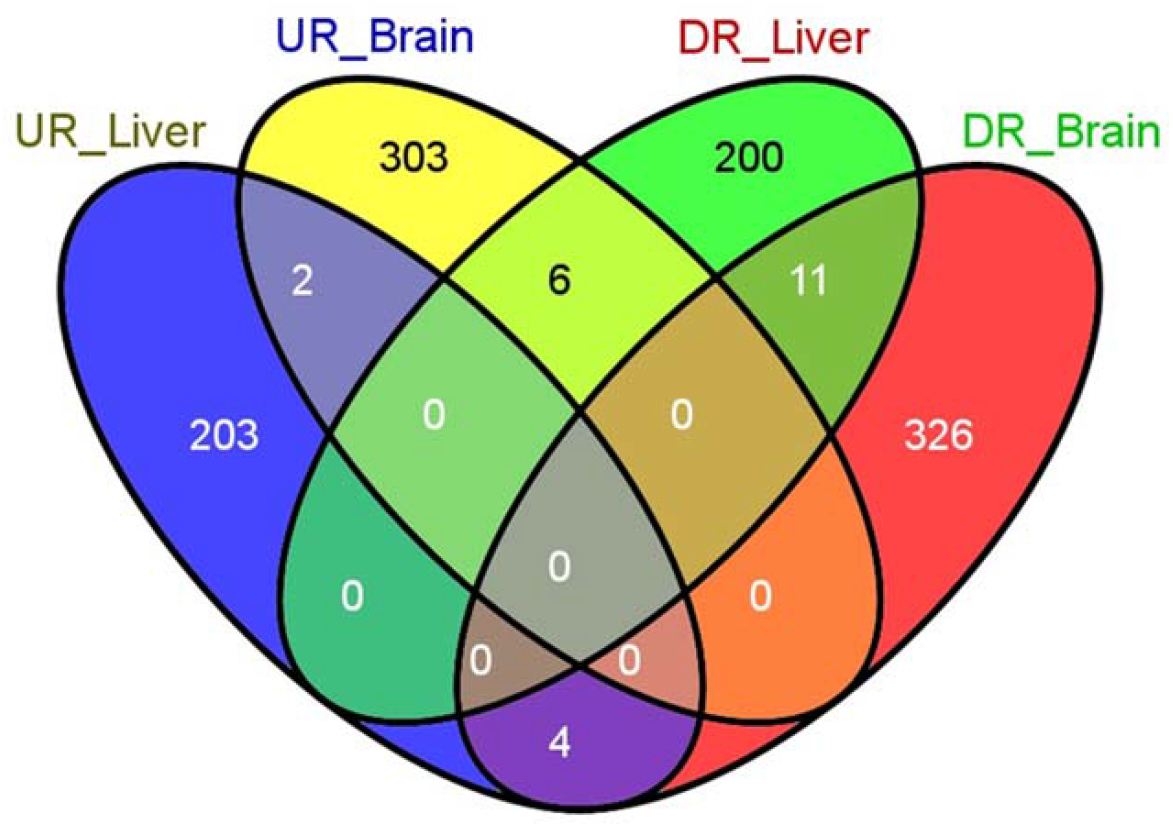
Venn diagram of the differentially expressed genes in the liver and brain after a rapid eye movement of sleep deprivation for 9 days in the rat. The Venn diagram shows overlapping genes of the UR_Liver: up-regulated liver; UR_Brain: up-regulated brain; DR_Liver: down-regulated liver; DR_Brain: down-regulated brain. Number in separate shaded panels reflects the genes typically affected in both tissues, in a similar or opposite direction.

Out of this pool, we found a set of genes that were commonly affected, either in the same or opposite direction, between the brain and the liver. For example, 4 of the 11 genes identified (**Supplementary Table, ST-3A**; namely ***WEE1 G2 Checkpoint Kinase (Wee1), Solute Carrier Family 2 Member 12 (Slc2a12), Harakiri, BCL2 Interacting Protein (Hrk)***, and ***Family With Sequence Similarity 110 Member B (Fam 110b)*** were negatively affected in both the brain and liver tissues (**Fig. 2**). Similarly, only 3 of the 6 genes identified (**Supplementary Table, ST-3B**; namely ***Hemoglobin Subunit Alpha 1 (Hba-a1)*** and ***Major urinary protein 5 (Mup5)*** were up-regulated in the brain and down-regulated in the liver (**Fig. 2**). In addition, we identified 3 genes (**Supplementary Table, ST-3C**; namely ***Histocompatibility 2, class II DR alpha (RT1-Da), Zinc Finger And BTB Domain Containing 6 (Zbtb6)***, and ***Transmembrane protein 106B (Tmem 106a)*** out of a total of 4 genes that were up-regulated in the liver and down-regulated in the brain (**Fig. 2**). In order to deepen our analysis, we moved forward with gene ontology and KEGG pathway analysis.

### 3.2. Gene Ontology analysis

Functional categories of genes that vary in their regulation between brain and liver upon REM sleep deprivation have been categorized. All processes and components were separated according to three main groups, namely biological processes, molecular functions, and cellular components (**Fig. 3-5**). In addition, we classified each group into two subcategories based on their direction of change (upregulation and downregulation), e.g. biological processes (**Fig.3, A-D**), molecular functions (**Fig. 4, A-D**) and cellular components (**Fig. 5, A-D**).

**Figure 3.**
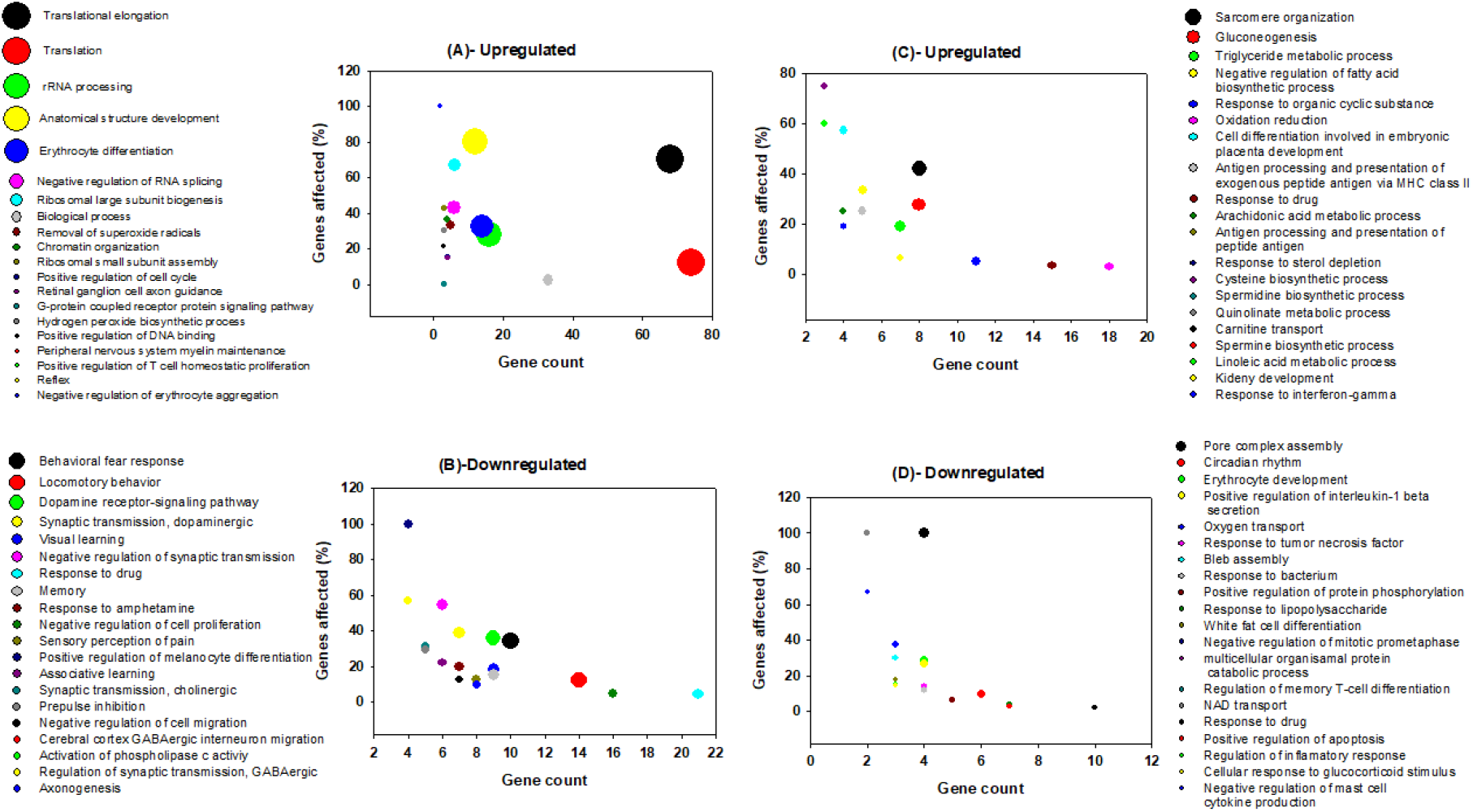
Graphical representation **of** GO terms from biological processes for genes which are up-regulated in the brain (A), down-regulated in the brain (B), up-regulated in the liver (C) and down-regulated in the liver (D) following rapid eye movements sleep deprivation in rats. The *x*-axis shows gene count, and the *y*-axis shows the percentage of genes affected for respective node. The bubble size represents the log transformed *p*-value [Y=−0.5*log(Y)] of the respective biological processes. A bigger bubble size indicates a more significantly affected a given process, and thus, a lower *p*-value. The top 20 terms are displayed for each category.

**Figure 4.**
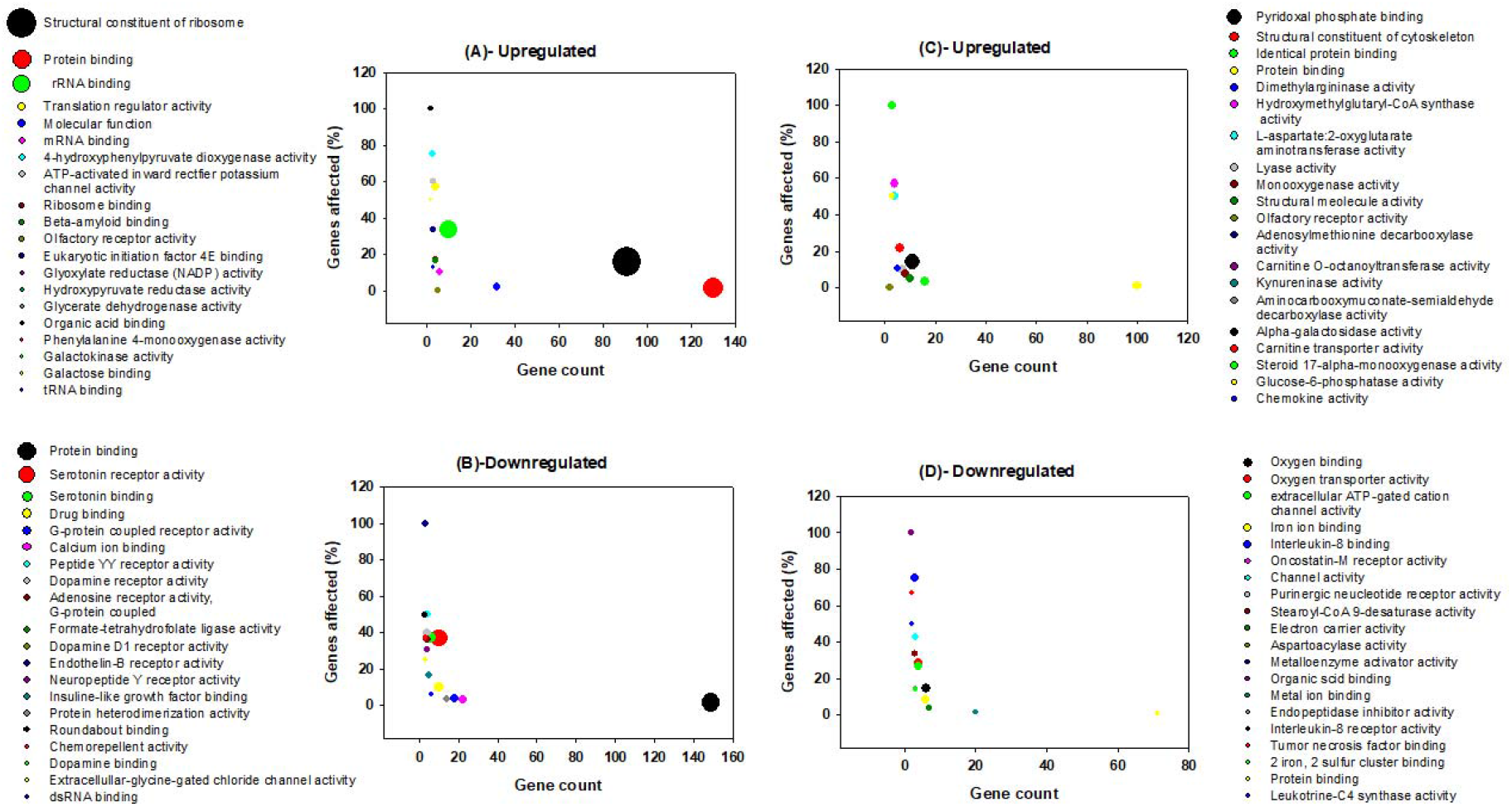
Graphical representation of molecular function terms for genes which are up-regulated in the brain (A), down-regulated in the brain (B), up-regulated in the liver (C) and down-regulated in the liver (D) following rapid eye movements sleep deprivation in rats. The *x*-axis shows gene count and the *y*-axis shows the percentage of genes affected. The bubble size represents the log transformed *p*-value [Y=−0.5*log(Y)] of the respective molecular function. A bigger bubble size indicates a more significantly affected a given process, and thus, a lower *p*-value. The top 20 terms are displayed for each category.

**Figure 5.**
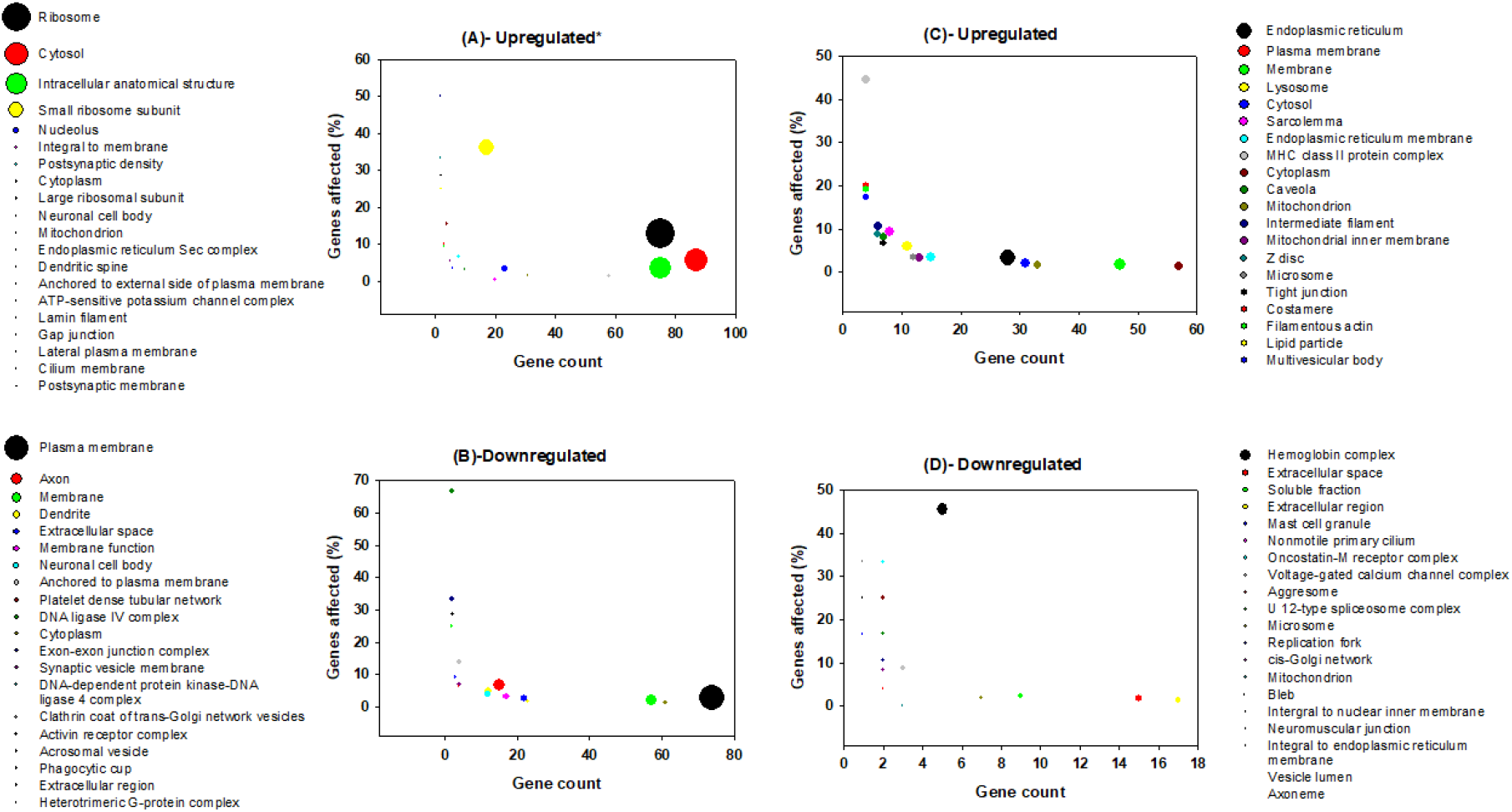
Graphical representation of cellular component terms for genes which are up-regulated in the brain (A), down-regulated in the brain (B), up-regulated in the liver (C) and down-regulated in the liver (D), following rapid eye movements sleep deprivation in rats. The *x*-axis shows gene count and the *y*-axis shows the percentage of genes affected. The bubble size represents the log transformed p-value [Y=−0.5*log(Y)] of the respective cellular component. To fit the bubble size, the *p*-value in 5A* was normalized with formula [Y=−0.5*log(Y)/2]. A bigger bubble size indicates a more significantly affected a given process, and thus, a lower *p*-value. The top 20 terms are displayed for each category.

Among the 208 significant GO terms of biological processes for genes which are upregulated in the brain, the top five are translational elongation, translation, rRNA processing, anatomical structure development and erythrocyte differentiation (**Fig. 3A**). Among the 77 significant GO terms of molecular functions for genes which are upregulated in the brain, the top five are structural components of ribosomes, protein binding, rRNA binding, translation regulator activity, and mRNA (**Fig. 4A**). Among the 57 significant GO terms of cellular components for genes which are upregulated in the brain, the top five include ribosomes, cytosol, intracellular anatomical structure, small ribosomal subunits, and nucleolus (**Fig. 5A**). REM sleep loss negatively affected 544 biological processes in the brain, of which the top five were behavioral fear response, locomotory behavior, dopamine receptor signaling pathway, dopaminergic synaptic transmission, and visual learning (**Fig. 3B**). A total of 140 significant molecular function terms were returned for genes which were negatively affected in the brain and the top five were protein binding, serotonin receptor activity, serotonin binding, drug binding and G-protein coupled receptor activity (**Fig. 4B**). A total of 57 cellular component terms were returned for genes which are negatively affected in the brain, of which the top five were plasma membrane, axon, membrane, dendrite, and extracellular space (**Fig. 5B**).

The top five of 355 significant biological processes terms for genes which were positively affected in the liver are Sarcomere organization, Gluconeogenesis, triglyceride metabolic process, negative regulation of fatty acid biosynthetic process, and response to an organic cyclic substance (*Fig. 3C*). The top five of 150 significant molecular function terms for genes which were positively affected in the liver are pyridoxal phosphate binding, structural constituents of the cytoskeleton, identical protein binding, protein binding, and dimethyl arginase activity (**Fig. 4C**). The top five of 64 significant cellular component terms for genes which were positively affected in the liver are endoplasmic reticulum, plasma membrane, membrane, lysosome, and cytosol (**Fig. 5C**). Pore complex assembly, circadian rhythm, erythrocyte development, positive regulation of interleukin-1 beta secretion, and oxygen transport were the top five of 219 significant biological processes terms (**Fig. 3D**) for genes which were downregulated in the liver. Oxygen binding, oxygen transport activity, extracellular ATP-gated cation channel activity, Iron-ion binding, and Interleukin-8 binding were the top five of 86 significant molecular function terms for genes which were downregulated in the liver (**Fig. 4D**). Hemoglobin complex, extracellular space, soluble fraction, extracellular region, and mast cell granules were the top five of 27 significant cellular component terms for genes which were downregulated in the liver (**Fig. 5D**).

### 3.3. Pathway analysis

KEGG analysis was used to evaluate the pathways affected by REM sleep loss in the brain and liver, and terms were plotted based on significance level (*p*<0.05), database count, and the number of genes affected by each pathway (node count). Shown are up- and down-regulated pathways in the brain (**Fig. 6**), up-regulated pathways in the liver (**Fig. 7A**), and down-regulated pathways (**Fig. 7B**). Pathways that were significantly upregulated in the brain included only ribosomes and olfactory transduction, while 11 were downregulated—of which the top five were neuroactive ligand-receptor interaction, axon guidance, calcium signaling pathway, olfactory transduction, and GAP junction (**Fig. 6**). The top five of 36 significantly upregulated liver pathways were glyceraldehyde metabolism, alanine and aspartate metabolism, cysteine metabolism, cell adhesion molecules, and glycine-serin & threonine metabolism (**Fig. 7A**), while just circadian rhythm, arachidonic acid metabolism, nitrogen metabolism, and retinol metabolism were downregulated (**Fig. 7B**).

**Figure 6.**
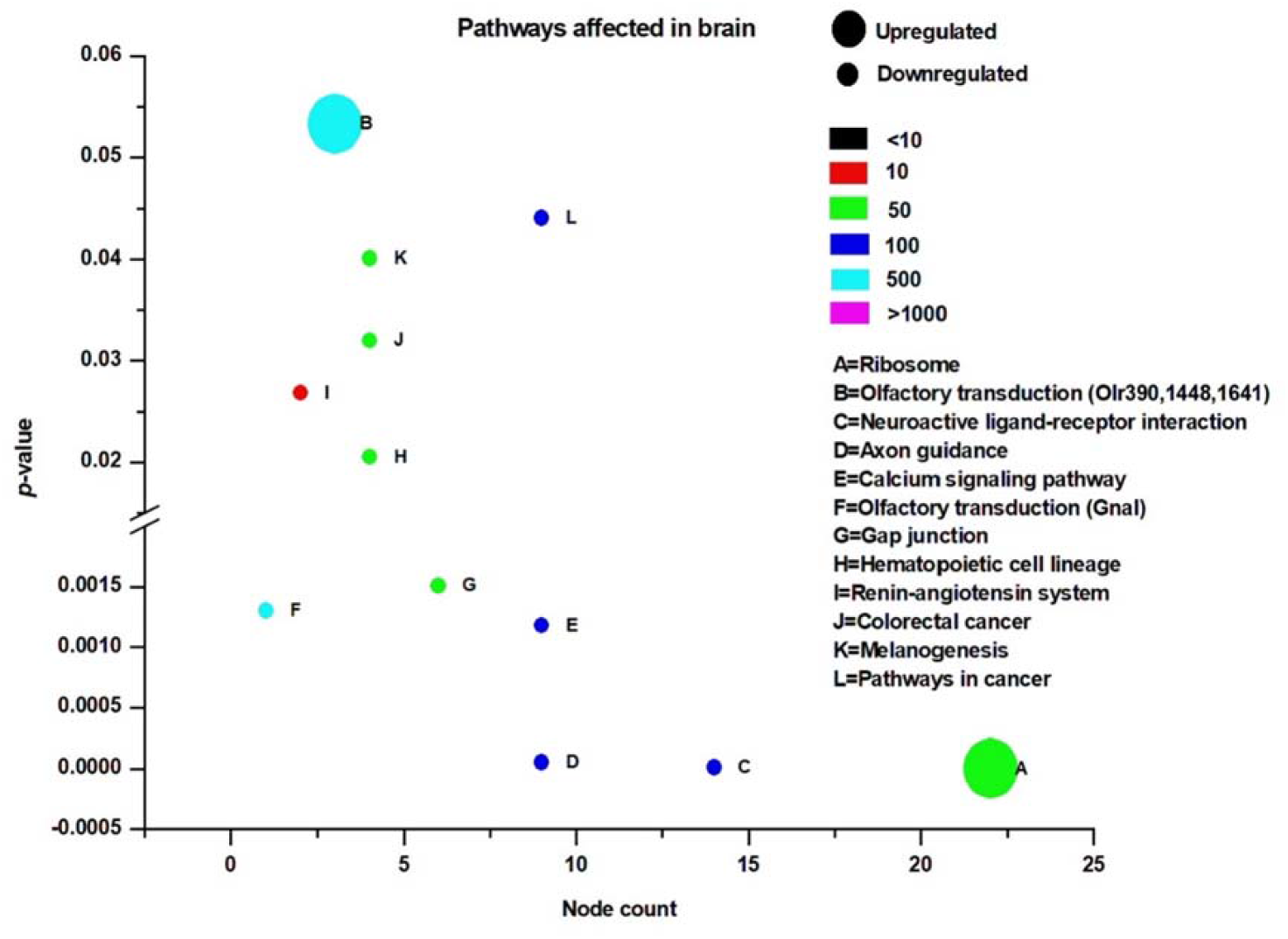
KEGG pathways affected by rapid eye movement sleep deprivation in rats in the brain. The *x*-axis depicts the number of nodes affected and the *y*-axis shows the *p*-value (*p*<0.05). Color coding indicates the total number of database nodes evaluated. The size of the circle indicates the direction of change.

**Figure 7.**
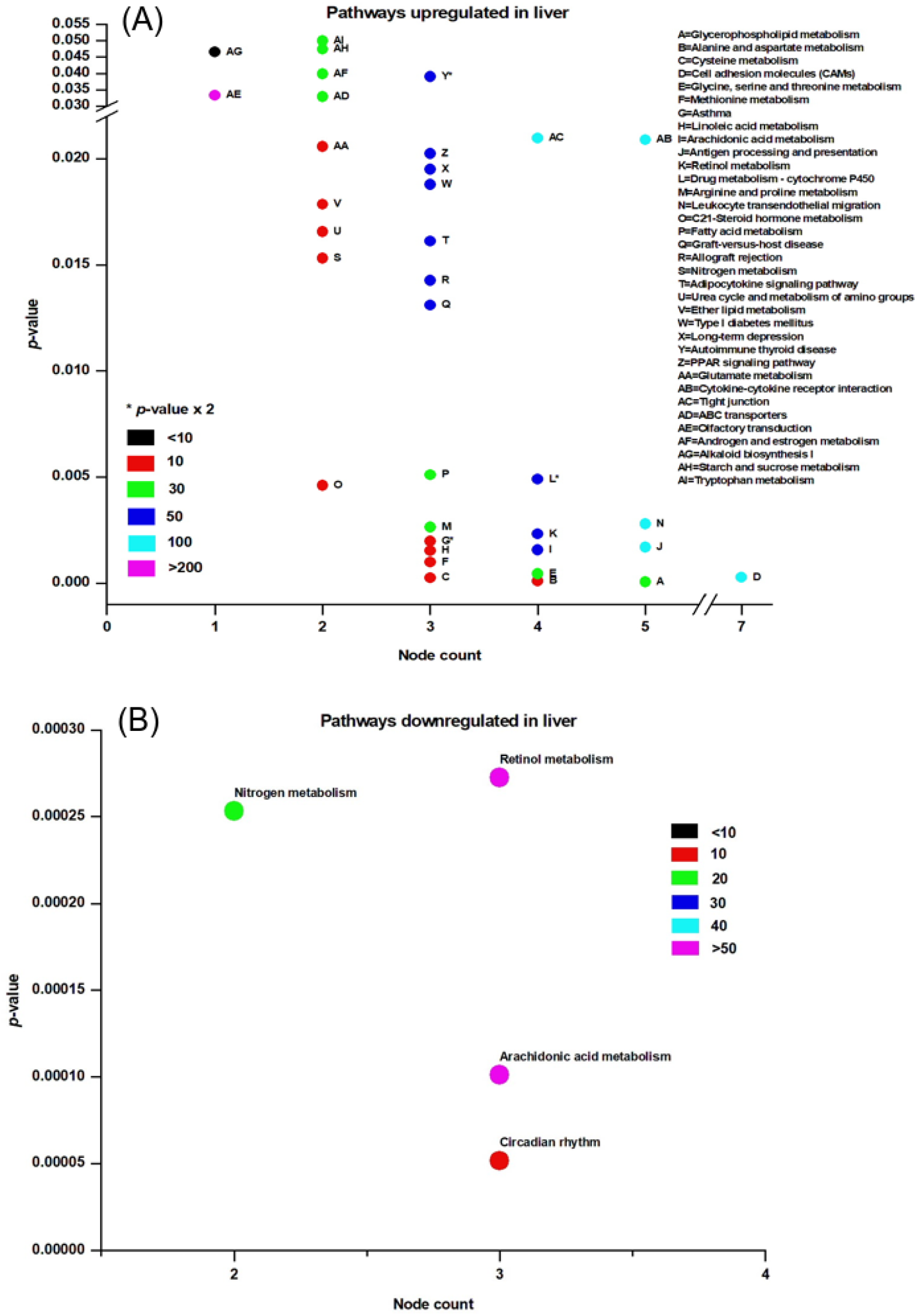
KEGG pathways affected by rapid eye movement sleep deprivation in rats in the liver. (A) Up-regulated, (B) Down-regulated. The *x*-axis depicts the number of nodes affected and the *y*-axis shows the *p*-value (*p*<0.05). Color coding reflects the total number of associated nodes.

We used a KEGG pathway maps (**Supplementary Figures, SF 1**-**4**) to visualize the components, proteins, and genes that were highly associated with the involved processes. Displayed are the subunits of ribosomes (**Supplementary Figure, SF-1**) and neuroactive legend-receptor interaction pathways (**Supplementary Figure, SF-2**), which were negatively and positive affected pathways, respectively, in the brain. Additionally, the cancer pathway (**Supplementary Figure, SF-3**) and glycerophospholipid metabolism pathway (**Supplementary Figure, SF-4**), which were negatively and positively affected, respectively, due to REM sleep loss.

## 4. Discussion

We sought to characterize the effects of prolonged REM sleep deprivation using gene expression data from the brain and liver of rats. In order to confirm our findings and validate our analyses, six differentially expressed genes were analysed using RT-PCR (*Fig.1*). We found that our study corroborates with previous microarray findings of sleep deprivation relating to effects on genes such as ***Egr1, Fos, Ptgs2***, several genes of Slc family, ***Hba-a1*** and ***Hbb*** [82,99–101]. Indeed, each of these six genes were found to be associated with sleep loss previously. Out of the hundreds of genes found to be significantly affected in the brain and liver, only a few genes were common between the tissues examined and their associated direction of change. Four genes, ***Wee1, slc2a12, Hrk***, and ***Fam110b*** (**Fig. 2, Supplementary Table, ST-3A**), were commonly downregulated in the tissues examined but none were commonly upregulated. Three genes, ***Hba-a1, Hba-a2 & Mup5***, were up-regulated in the brain and down-regulated in the liver and associates with GO terms drug transport, oxidoreductase activity, heme binding, fatty acid biosynthesis processes, and catalytic activity (**Fig. 2, Supplementary Table, ST-3B**). Genes that were found to be up-regulated in the liver, and at the same time, down-regulated in the brain (***RT1Da, Zbtb6, and Tmem 106b***) are associated with the GO term stimulus response (**Fig. 2, Supplementary Table, ST-3C**).

Several of the aforementioned genes which were commonly associated between brain and liver tissue, and any combination of direction of change, were found in previous literature regarding sleep and REM. Several genes of solute carrier (Slc) family (**Supplementary Table, ST-4**) were up- and down-regulated in the brain and liver, respectively, except ***slc2a12*** which was downregulated in both the brain and liver. Previously, genes of the ***slc*** family were reported to be associated with glucose homeostasis and ***slc17a8*** is down-regulated in Tinaja cave fish in response to sleep deprivation [102]. ***Slc38a5a*** is upregulated in response to sleep deprivation when glucose levels drops and circulating amino acid levels increases [103]. Recently, ***Hrk*** gene was found to be upregulated in mice after sleep deprivation, which is opposite to that of our findings and may be a result of differential expression between organisms or sleep-loss in general, as compared to only REM deprivation [104]. GO term analysis of molecular/biological functions associated with ***Hrk*** returned the terms protein tyrosine kinase activity, carbohydrate transmembrane transport activity, apoptosis regulation, and Bleb assembly (**Supplementary Table, ST-3A**). Previous studies and recent pre-prints support that REM sleep deprivation results in the apoptotic death of neuronal and hepatocytic cells [44–46]. Induction of ***Hba-a1*** gene in brain may cause cerebral hypoxia-like condition in the brain after REM loss as a result of cerebral hypoxemia and obstructive sleep apnea, and reduce hemoglobin denaturation [105,106]. A recent study on sleep restriction showed that there is an increase in free fatty acids in healthy men, which led us to speculate that REM sleep deprivation can affect genes such as ***Mup5***, which our findings demonstrated an association with the term fatty acid biosynthetic processes and was differentially expressed as a result of REM sleep deprivation [58]. Similarly, ***Zbtb6*** is a homologous gene that codes for the BTB domain of zinc finger protein in mammals and ***Tmem 106b*** returned several GO terms, which include protein binding, dendrite morphogenesis, and lysosomal transport [107,108]. A recent study showed that ***Tmem 106b*** is associated with dementia, which is caused by faulty regulation of microRNAs [109]. Overall, our study provides a list of genes affected across different tissue of body and further commonly affected in same or different directions which would be interesting to explore in future.

Many of the GO terms in our findings indicated the presence of various phenomena associated the synapse, and more specifically, synaptic potentiation (**Fig 3-5**). Previous analyses demonstrated that several genes, ***Arc, Bdnf, Camk4, Creb1, Egr1, Fos, Nr4a1, Ppp2ca***, and ***Ppp2r2d*** are associated with the GO term, potentiation of synaptic plasticity, which largely supports the ‘synaptic homeostasis theory’ [55,58]. Indeed, we found ***Fos*** and ***Egr1*** is significantly downregulated in our study in both brain and liver. In addition, several other genes (list not shown) of our study that were non-significantly upregulated/downregulated in the liver/brain, respectively are associated with GO terms related to synaptic plasticity, like positive regulation of long-term neuronal synaptic plasticity, regulation of neuronal synaptic plasticity, synaptic vesicle endocytosis, and neuromuscular synaptic transmission. Previously sleep loss has shown to be involved in the upregulation of genes associated with synaptic plasticity [110–112], however, many of its associated GO terms (**Supplementary Table, ST-5**) were a result of genes which were downregulated in the brain of REM sleep loss rats in our study. The regulation of synaptic plasticity during sleep and learning is essential [113] and loss of sleep was found to be associated with negative impact on glial signaling pathway important for synaptic plasticity [25,114–116].

REM sleep deprivation is found associated with modification of expression of long-term potentiation in visual cortex of immature rats [117] and we report upregulation of structural constituents of ribosomes, translation regulation activity, while dopamine receptor-signaling pathway, dopaminergic, cholinergic, GABAergic regulation of synaptic transmission, serotonin binding, and receptor activity were downregulated in brain (**Fig. 3B & 4B**). The dopamine receptor-signaling pathway regulating sleep, learning, and its plasticity are well known [84,85]. Sleep disorders and sleep deprivation have been correlated with dopaminergic, cholinergic, and GABAergic regulation of synaptic transmission, each of which were terms that were significantly enriched for genes that were downregulated in our study [86–90]. These observations support our hypothesis that REM sleep loss negatively affect the gens and processes related to synaptic homeostasis in brain.

Processes and pathway in the liver following REM sleep deprivation are largely associated with metabolism and the immune system. Many metabolic process and cellular metabolic processes such as gluconeogenesis, triglyceride metabolic process, negative regulation of fatty acid biosynthetic process, oxidation reduction, and arachidonic acid metabolic process were upregulated in liver in response to REM loss. Whole body energy expenditure decreases by 15-35 percent, With the lowest expenditure during slow-wave sleep and marginally higher during REM sleep [118] and sleep restriction involves reduced muscle glucose uptake, elevated blood glucose production, and pancreatic β-cell dysfunction [119,120]. A increasing body of evidence indicates that Obstructive Sleep Apnea Syndrome is associated with a variety of metabolic alterations, such as dyslipidemia, insulin resistance, glucose intolerance [121]. REM sleep impairs glucose metabolism that is involved in intermittent hypoxemia [122]. An upregulation of gluconeogenesis may serve as a mechanism to compensate for hypoxemia due to prolonged REM loss. The GO terms related to homeostatic processes like cholesterol homeostasis, nitric oxide homeostasis, fatty acid homeostasis, retina homeostasis, cytosolic calcium ion homeostasis are associated with genes which were upregulated in the liver, while T cell homeostasis and other processes associated with the immune system were downregulated (**Supplementary Table, ST-6**). The immune functions of sleep and associated diseases have been studied [123,124], as well as and evidence that the immune system is compromised from lack of sleep. [125]. The body of previous evidence and our results support our hypothesis, that while REM sleep loss is associated with synaptic potentiation and maintenance, its affects in the liver are more so related to metabolism and immune response to infections.

REM sleep loss negatively affects several genes linked to neuroactive ligand-receptor interaction pathways in brain, primarily gamma-Aminobutyric acid, Human Thrombin receptor, and associated receptor signaling dopamine (**Supplementary Figure, SF-2**). A recent review of sleep and protein-dependent synaptic plasticity, indicated that sleep deprivation impairs many of the related biological and physiological processes [126]. We have found that many of the pathways in the liver which have been upregulated are linked to metabolism, immunity, and depression (**Fig. 7A**). On the other hand, only a few downregulated pathways in the liver have been established, which include nitrogen metabolism and circadian rhythm (**Fig. 7B**). The findings further support our secondary hypothesis that REM sleep loss affects the processes and pathways related to synaptic potentiation and learning and memory (**Fig 8A**) and processes related to homeostasis and immunity in liver (**Fig. 8 A-C**).

**Figure 8:**
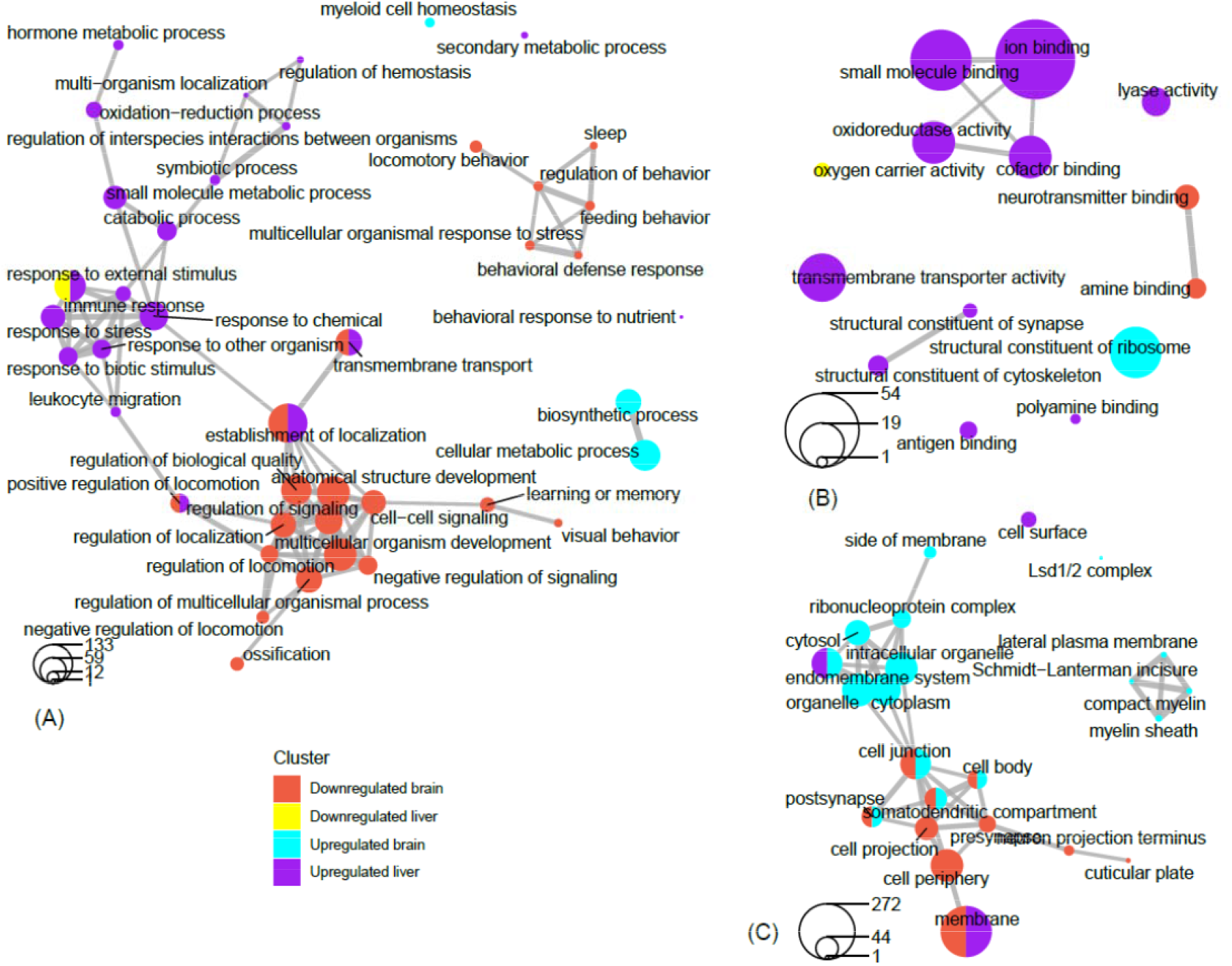
Network view of GO term association. Network plots of the top filtered GO terms, depicting the degree of connectivity within and between terms of enriched genes which are upregulated in the brain, upregulated in the liver, downregulated in the brain, and downregulated in the liver. The circles-legend at the bottom of each left-hand corner indicates the number of genes that are enriched for a given term. Connecting lines indicate a significant degree of semantic similarity between terms. Biological process (A), cellular component (B), and molecular function (C). GO terms were filtered (level = 3) to reduce redundancy and capture major categorical themes prior to visualization of connectivity in network plots, which were designed in R using cluster Profiler package. Plots of filtered GO terms contained the top 20 significant categories, respectively, per subject cluster.

Network analysis of filtered GO terms allowed for visualization of major themes and the connectivity of processes across brain and liver tissue in rats deprived of REM (**Fig. 8**). Several biological processes, like positive regulation of locomotion, establishment of localization, and transmembrane transport were terms that were significantly enriched for genes that were both downregulated in brain and upregulated in liver. Interestingly, response to external stimuli genes were found as both positively and negatively affected in the liver, indicating the up- and downregulation of separate sets of genes associated with this term (**Fig. 8A**). There was no connectivity between terms in the molecular function category, however, terms associated with metabolism and transport, like oxidoreductase activity, small molecule binding, iron binding and cofactor binding, were each upregulated in liver (**Fig. 8B**). Networking of terms in the cellular component category returned GO terms, cell junction, cell body, post synapse, and somatodendritic compartment, that were up- and downregulated in the in the brain and liver, respectively (**Fig. 8C**). To summarize a major theme, some processes which were mainly upregulated in liver were also downregulated in the brain as a result of REM sleep loss. One possible explanation for this is that REM sleep loss influences processes linked to fear response of the brain and locomotive activity related to the peripheral circadian clock, hemoglobin level, and transport of oxygen throughout the liver. The evidence suggests that the genes and processes involved are highly contrasted between the brain and liver, however, some processes may be connected across major organs in response to REM sleep loss and should be investigated in the future.

We further explored the common processes related to general interest like oxidative-stress, cancer, and cell-death. Processes related to reactive oxygen species metabolic and oxidative stress e.g., positive regulation of oxygen and reactive oxygen species metabolic process, response to oxidative stress were positively affected while, cellular response to reactive oxygen species, oxygen and reactive oxygen species metabolic processes negatively got affected in brain. Previous studies have shown that there are transcript level variations in many genes involved in the regulation of reactive oxygen species (ROS), including heme oxygenase, superoxide dismutase, and catalase, in patients with obstructive sleep apnea [127]. Similarly, REM sleep is recently found associated with acute phase response and ROS stress in liver [47,49]. REM sleep loss also affected several genes such as ***prostaglandin-endoperoxide synthase (Ptgs2), B-cell lymphoma 2 (Bcl-2), Proto-Oncogene, Tyrosine Kinase receptor (Kit), KRAS Proto-Oncogene (K-Ras)*** and ***Fos Proto-Oncogene (Fos)*** which are marked in cancer pathways (**Supplementary Figure, SF-3**). A number of recent studies have shown that sleep dysfunction/loss and cancer processes are very related [128–134]. However, some emerging evidence also suggests that sleep loss/insomnia prior to the onset of cancer is independently associated with cancer risk [129,133,135,136]. ***Ptgs2***, an enzyme, plays a key role in various pathological processes by catalyzing conversion of arachidonic acid to prostaglandins [137].

Studies have shown that overexpression of ***Ptgs2*** is associated with angiogenesis, metastases, and immunosuppression [75,76]. ***Pgst2*** is also found to be associated with the chemoresistance of some malignant tumors, including liver, pancreatic, lung, esophageal, and gastric cancers [77–79]. Inhibition of ***Ptgs2*** effectively increased the sensitivity of tumors to drugs [138]. Similarly, ***Bcl-2, Kit, K-Ras***, and ***Fos*** genes have been found associated with cancer [139–142]. These genes play an important role in the sleep-wake cycle regulation and are shown to be correlated with sleep [44,45,143–145]. At the same time, glycerophospholipid metabolism pathway was found significantly upregulated in liver (**Supplementary Figure, SF-4**). These include the genes ***Phospholipase, PLa2g, Phosphatidylcholine 2-Acylhydrolase 12A Pla2g12a, Glycerol-3-Phosphate Dehydrogenase 2, Gpd2, CDP-Diacylglycerol Synthase 2, Cds2, and Phospholipid Phosphatase 2, Plpp2***. The PLa2g associates with neurodegeneration and elevated mitochondrial lipid peroxidation and dysfunction [146–148]. The ***PLa2g*** is further found positively associated with sleep loss and psoriasis in humans [149,150]. Similarly, ***Gpd2*** gene is found associated with intellectual disability in humans [151] and positively affected due to circadian desynchrony in mouse [152]. The chronic sleep deprivation in rats affected the protein profile of ***Gpd2*** in hypothalamic astrocytes [101] The functional aspect of other genes affected (e.g., ***Pla2g12a, Cds2*** and ***Plpp2***) is lacking and needs further exploration. These findings further support the idea of REM sleep related to restorative functions against diseases and oxidative stress.

Many KEGG pathways were associated with genes that were either significantly up- or downregulated (**Fig. 6 and Fig. 7A & 7B**) in the brain or liver as a result of REM sleep loss. The KEGG pathway map (**Supplementary Figures, SF 1-4**) demonstrated that many of the genes for ribosomal proteins that are involved in protein synthesis processes were upregulated in the brain by REM sleep loss (**Supplementary Figure, SF-1**). Indeed, research has shown that long-term sleep loss has been found to control several genes in the brain that are linked to DNA binding/regulation of transcription, immunoglobulin synthesis, and stress response [56,91]. Contrary to the notion that ***Homer-1a*** is a key brain molecule in response to sleep loss in mice, no effect on gene expression of Homer gene was observed in our study, which suggests that its regulation is modulated during other stages of sleep or an organism-specific phenomenon [82]. The results underscore the complexity of sleep loss and its associated consequences, and requires sleep phase-, species-, and/or tissue-specific considerations rather than overarching, vague generalizations to deeply understand the phenomenon.

Additionally, REM sleep loss negatively affected several genes linked to neuroactive ligand-receptor interaction pathways in brain, primarily related to gamma-Aminobutyric acid, Human Thrombin receptor, and associated receptor signaling dopamine (**Supplementary Figure, SF-2**). A recent review of sleep and protein-dependent synaptic plasticity, indicated that sleep deprivation impairs many of the related biological and physiological processes [126]. We have found that many of the pathways in the liver which have been upregulated are linked to metabolism, immunity, and depression (**Fig. 7A**). On the other hand, only a few downregulated pathways in the liver have been established, which include nitrogen metabolism and circadian rhythm (**Fig. 7B**). The findings further support our secondary hypothesis that REM sleep loss affects the processes and pathways related to synaptic potentiation and learning and memory (**Fig 8A**) and processes related to homeostasis and immunity in liver (**Fig. 8 A-C**).

Findings across studies are inconsistent in regard to REM sleep deprivation, and locomotor behavior and pain tolerance in rodents. Several studies have shown that REM sleep loss induces locomotor activity [93,153,154], while others have shown decreased locomotor activity [155]. The lack of consistent explanation could be related to procedural changes in the methods of a given study, such as the degree of REM sleep loss. Few studies have used multiple pots compared to our classic single flower pot method for deprivation, and other studies have implemented less total time for deprivation (72-96 hrs.) compared to ours, which was ~216 hrs. Recent research supports that REM sleep deprivation can affect locomotor activity in rats in an inverted-U manner [156,157]. A widely accepted view in the scientific community is that sleep deprivation decreases pain tolerance and increases the transmission of pain in multiple chronic pain conditions [158–163]. There is a conflict between reports on the sensory perception of pain [164,165] which was negatively affected in our study (**Fig. 3B**) and few studies indicated only total sleep deprivation raises the intensity of pain rather than REM sleep deprivation [166,167]. Nonetheless, selective REM sleep deprivation is correlated with enhanced placebo analgesia effects [168]. Similarly, consistency exists between REM sleep loss and its association with the perception of pain [170]. Perhaps sleep in general and short-term REM sleep deprivation lower the pain threshold, while long-term sleep deprivation increases the pain threshold. REM sleep deprivation and pain is significantly correlated with environmental conditions (e.g. dry or wet conditions), with pain sensitivity enhanced in dry test conditions but no different in wet conditions. [171]. This suggests that further work is needed to understand deeply the relationship between experience of pain and lack of sleep. Furthermore, a recent microarray analysis shows that ***Hspa5*** gene expression increases not only in the brain but also in the liver as sleep deprivation increases [82]. Our analysis did not return any genes which were commonly upregulated in both the brain and liver, however, this may simply be due to differences between species and many of the studies related to Hspa5 and sleep-wakefulness involved mice and drosophila [13,55,169]. Genes such as, ***Wee1, Slc2a12, Hrk***, and ***Fam110b*** were commonly downregulated in both the brain and liver. Currently, however, there is consensus about the relationship between expression of genes associated with locomotor behavior and pain tolerance, an area that is open for future research.

Our approach to GO term and KEGG pathway analysis is quite relevant in the current era of genomics and sequencing, but it also involves discrepancies in gene function across organisms, distributed biases and biases linked to positive and negative annotations (more information in 56–58). Like GO term analysis, KEGG analysis also has its limitations, apart from reducing the complexity of the data and helping to increase the explanatory power. One of the key disadvantages of KEGG is the independent consideration of pathways, even though crosses and overlaps occur in the natural system [177–180]. Therefore, the findings of our current study involving REM sleep deprivation affecting brain and liver should be taken as case study. The present study also provides as data set for future studies to compare the effect of RMS sleep loss across organs. We have only few microarray studies to understand effect of molecular signatures of diseases, effect of sleep deprivation, and disorders and looking forward to our study as one [127,181–186]. We further need more research related to total sleep or REM sleep loss to determine stage and tissue-specific effects on body in order to understand particular effects and to evaluate the influence of sleep loss and in effects of sleep related disorders.

## 5. Conclusion

Microarray analysis of brain and liver tissue in rats found that many of the physiological processes and the genes involved in the pathways are regulated differently in the two organs body as a consequence of REM sleep loss, which also supported our hypothesis that REM sleep is crucial for proper metabolism and immune function in the liver synaptic potentiation in the brain. Our findings underscore the idea that the brain have shown to be more receptive to processes such as synaptic potentiation, learning and memory, oxidative stress, and circadian rhythms in response to REM sleep loss. On the other hand, the function of the liver is more so related to processes like protein synthesis, stress balance, and detoxification. The study provides a fundamental platform for visualizing the effects of REM sleep loss across the brain and liver and future studies should address the underlying dynamics of REM sleep deprivation throughout the body.

## Acknowledgments

This research was carried out and funded by a laboratory grant from the SKK Laboratory at the School of Biotechnology, Jawaharlal Nehru University, New Delhi, India. AP received funding from the Research Fellowship of the University Grants Commission (2008-2012), India and afterward supported by planning and budgeting committee (PBC) outstanding postdoctoral fellowship of Israel to Chinese and Indian citizens (2014-2017) and project postdoc fellowship from Prof. Guy Bloch at The Hebrew University of Jerusalem, Israel.

## Conflicts of Interest

The authors have no conflict of interest and all funding and scientific contributions are fully recognized.

---

**Supplementary Figure 1.**
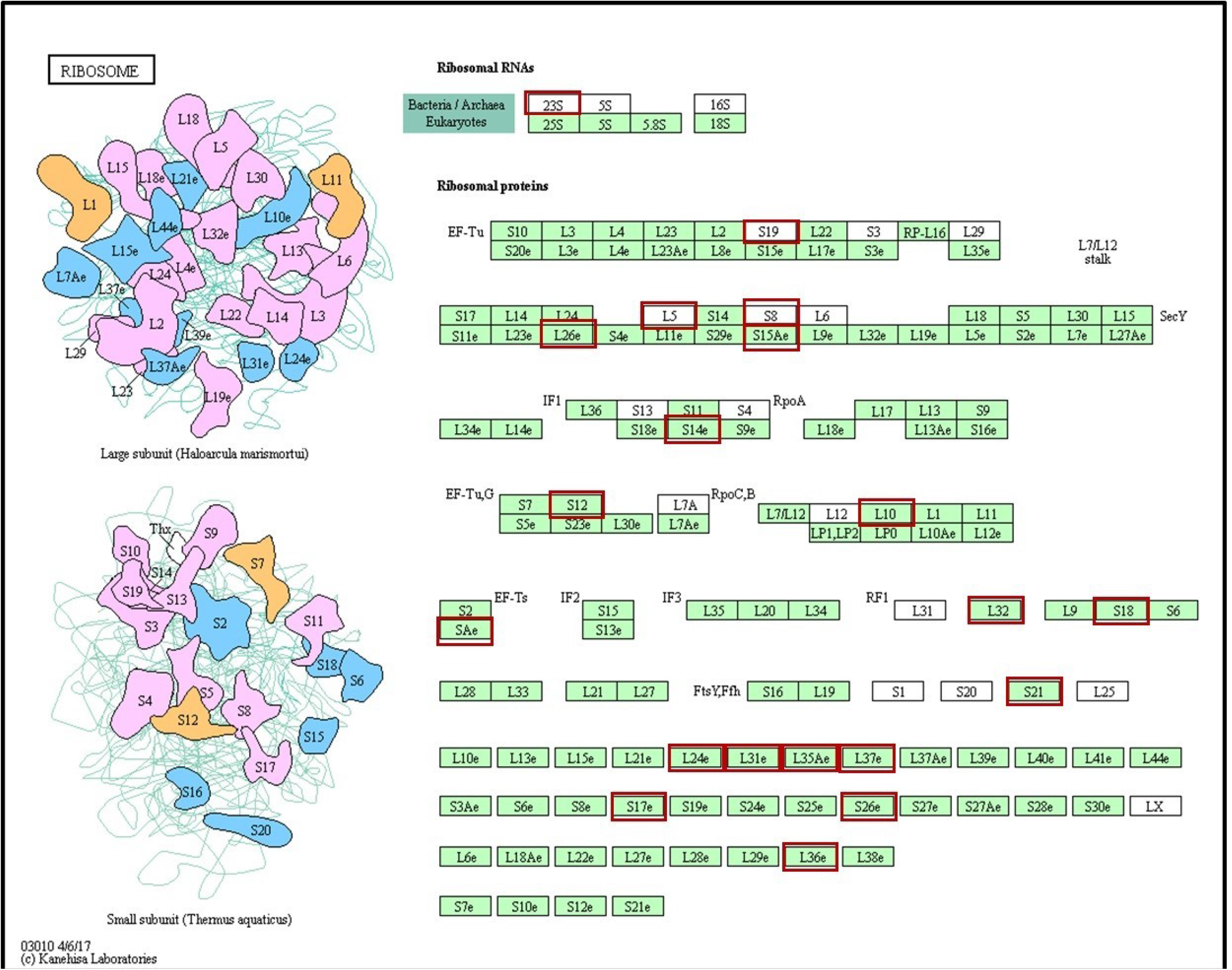

**Supplementary Figure 2.**
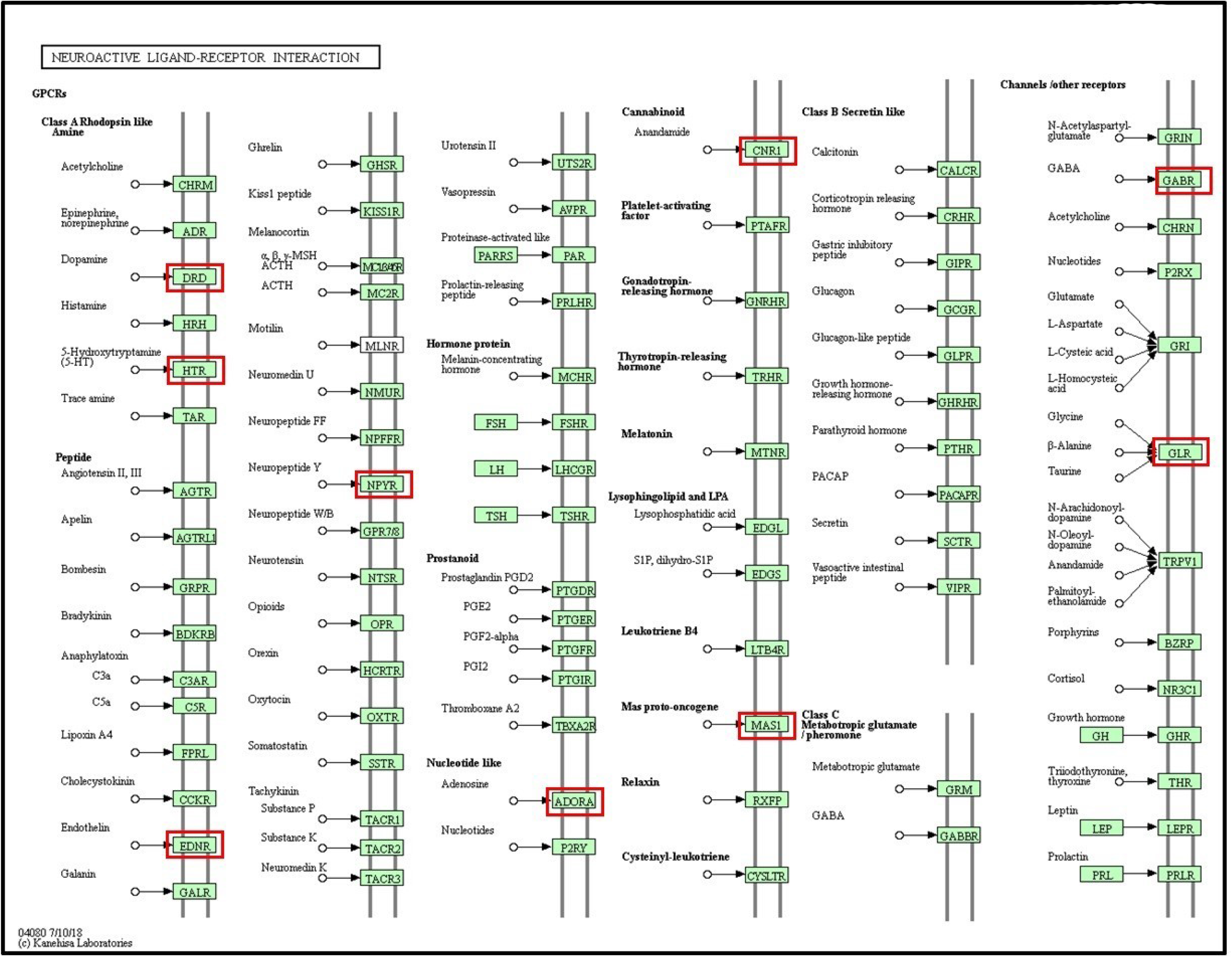

**Supplementary Figure 3.**
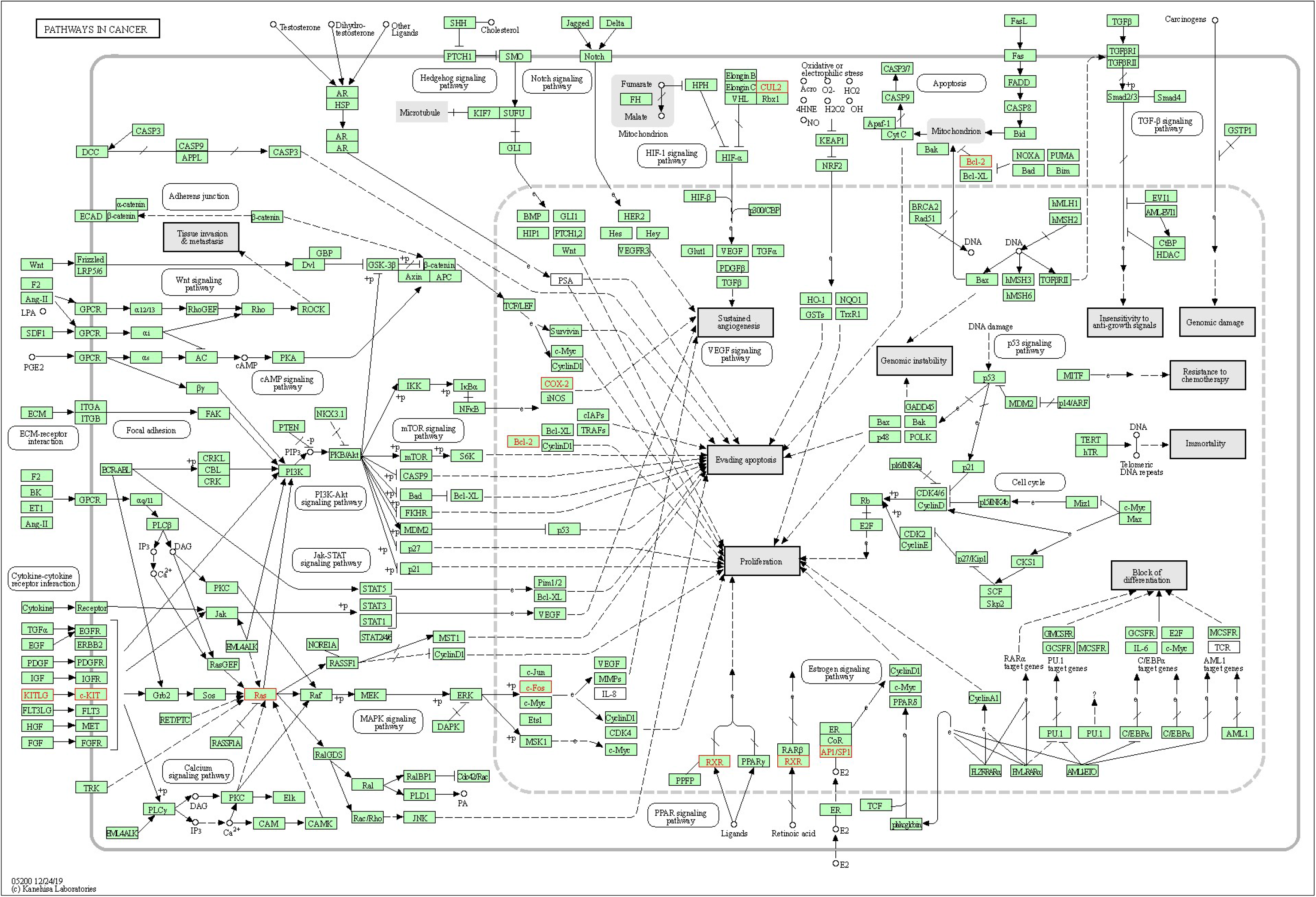

**Supplementary Figure 4.**
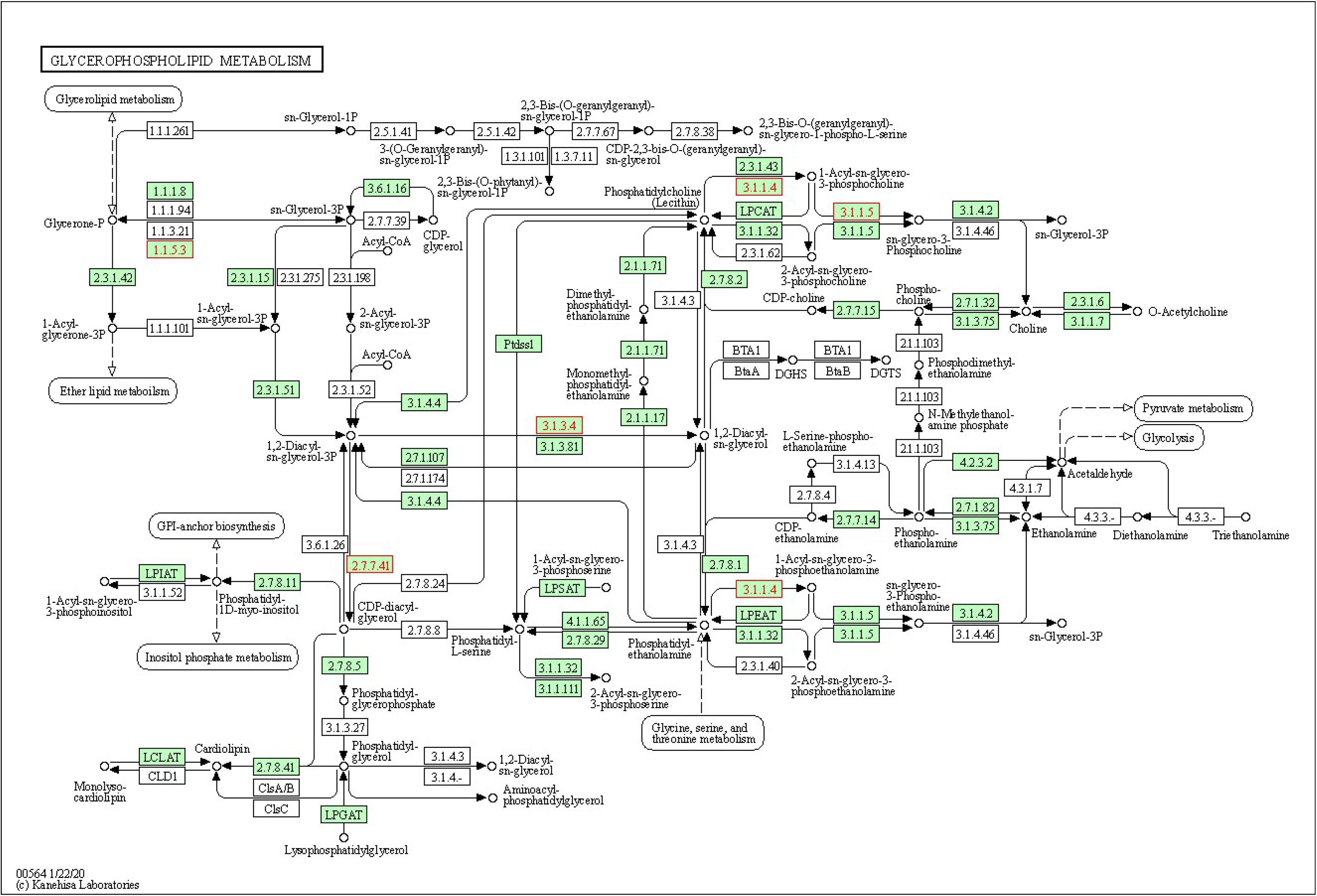

## Supplementary materials

**Supplementary table (ST1):**
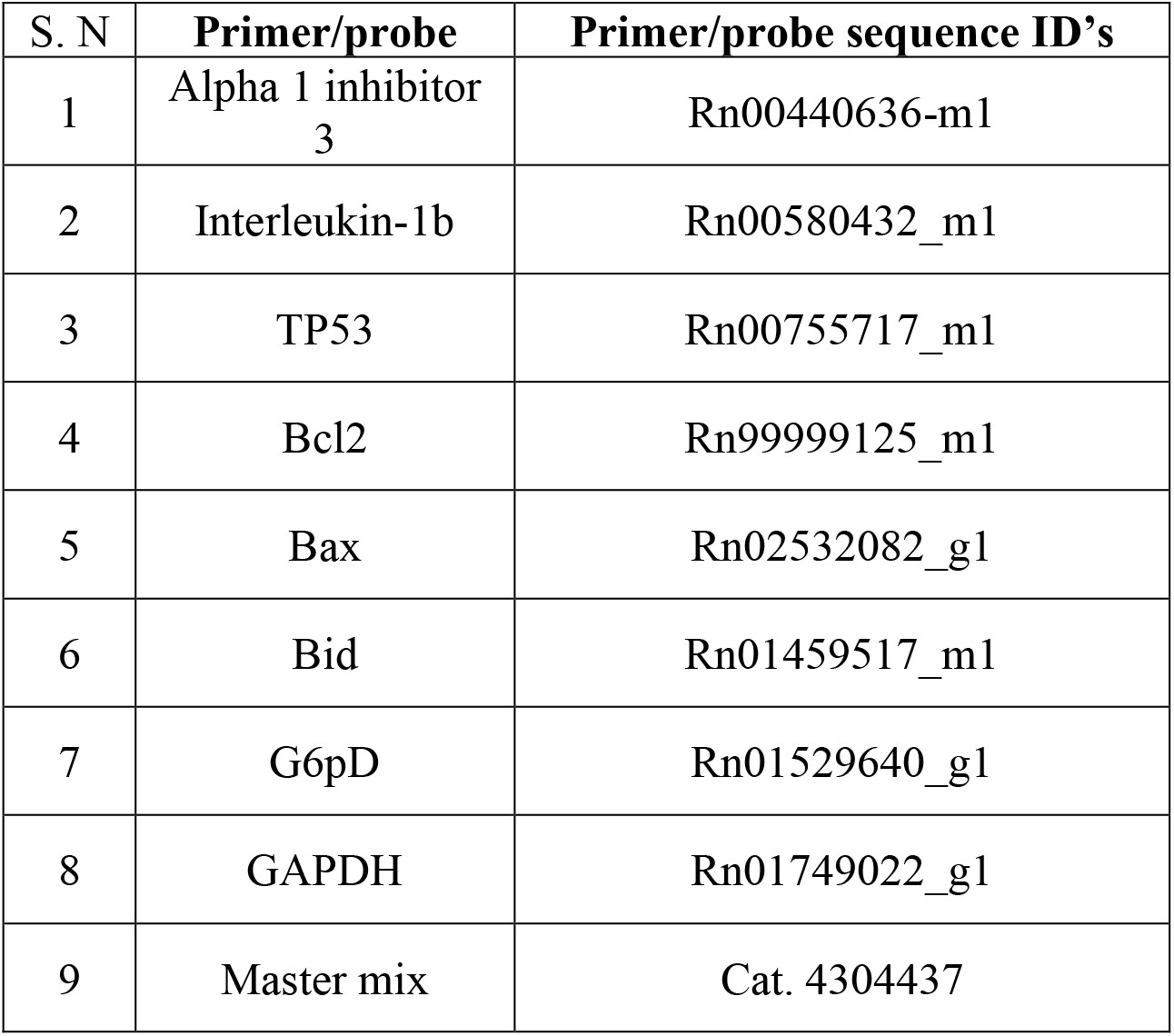
List of real time PCR gene primers used in microarray validation experiments.

**Supplementary table (ST2A):**
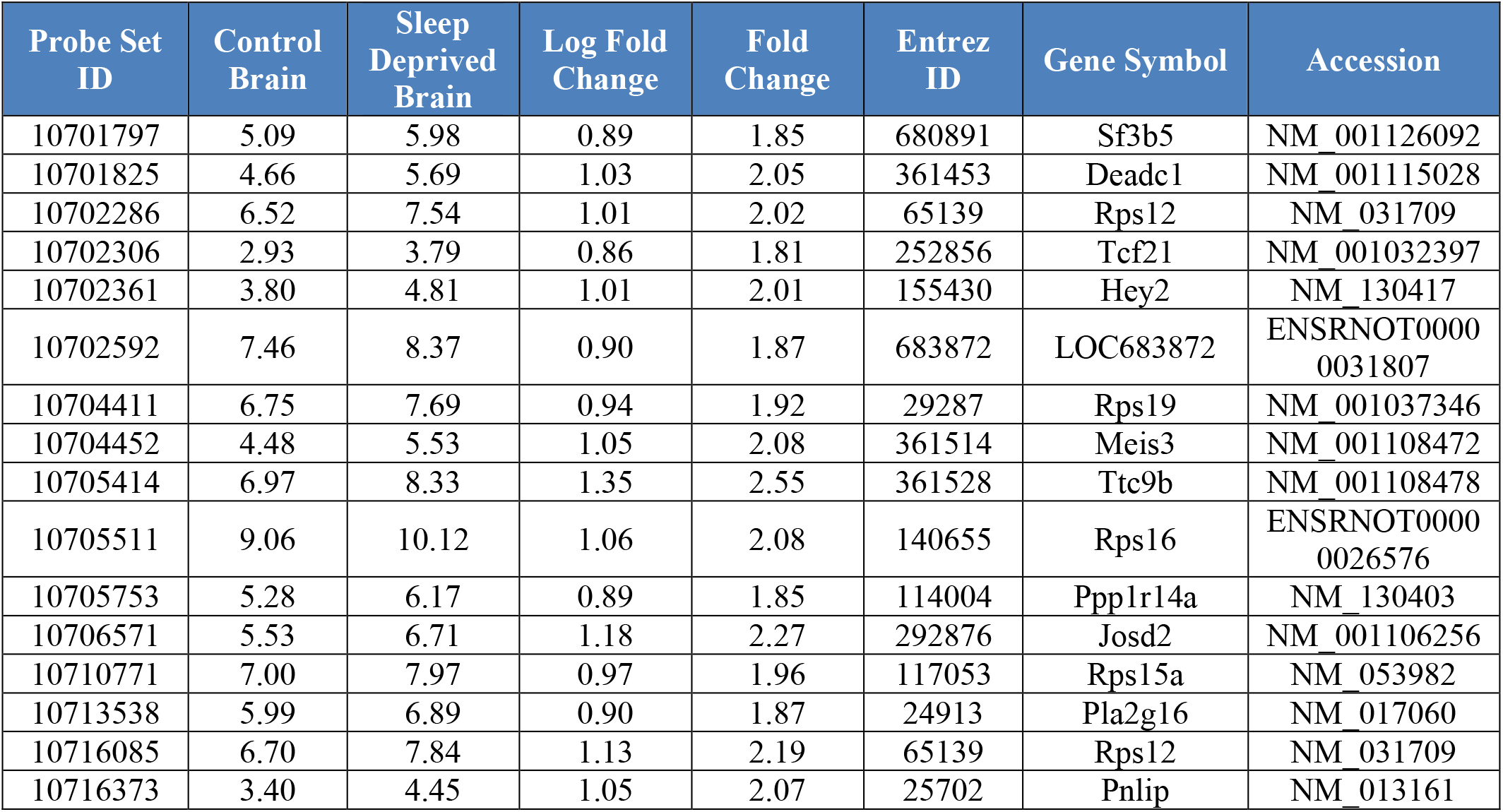
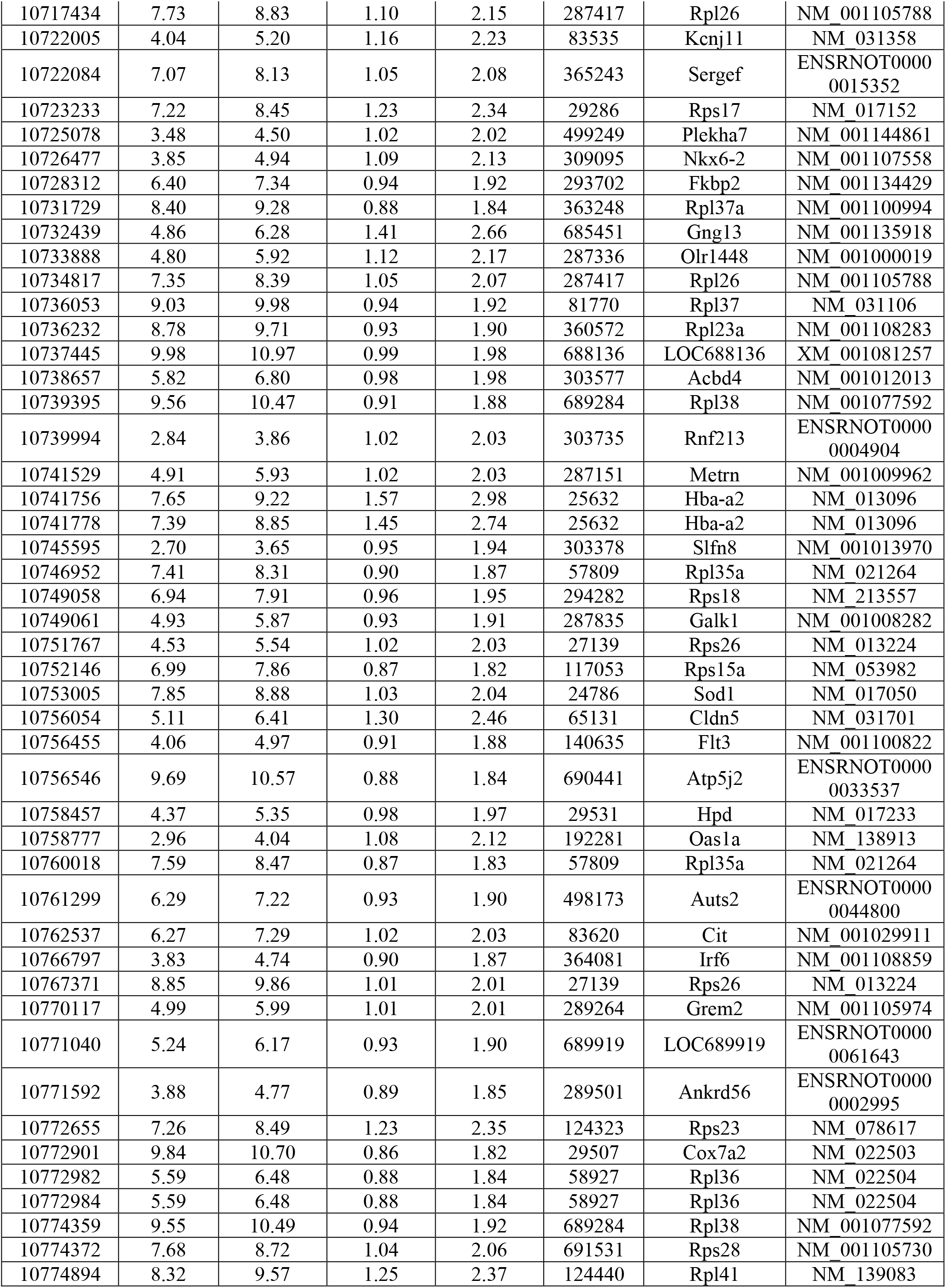
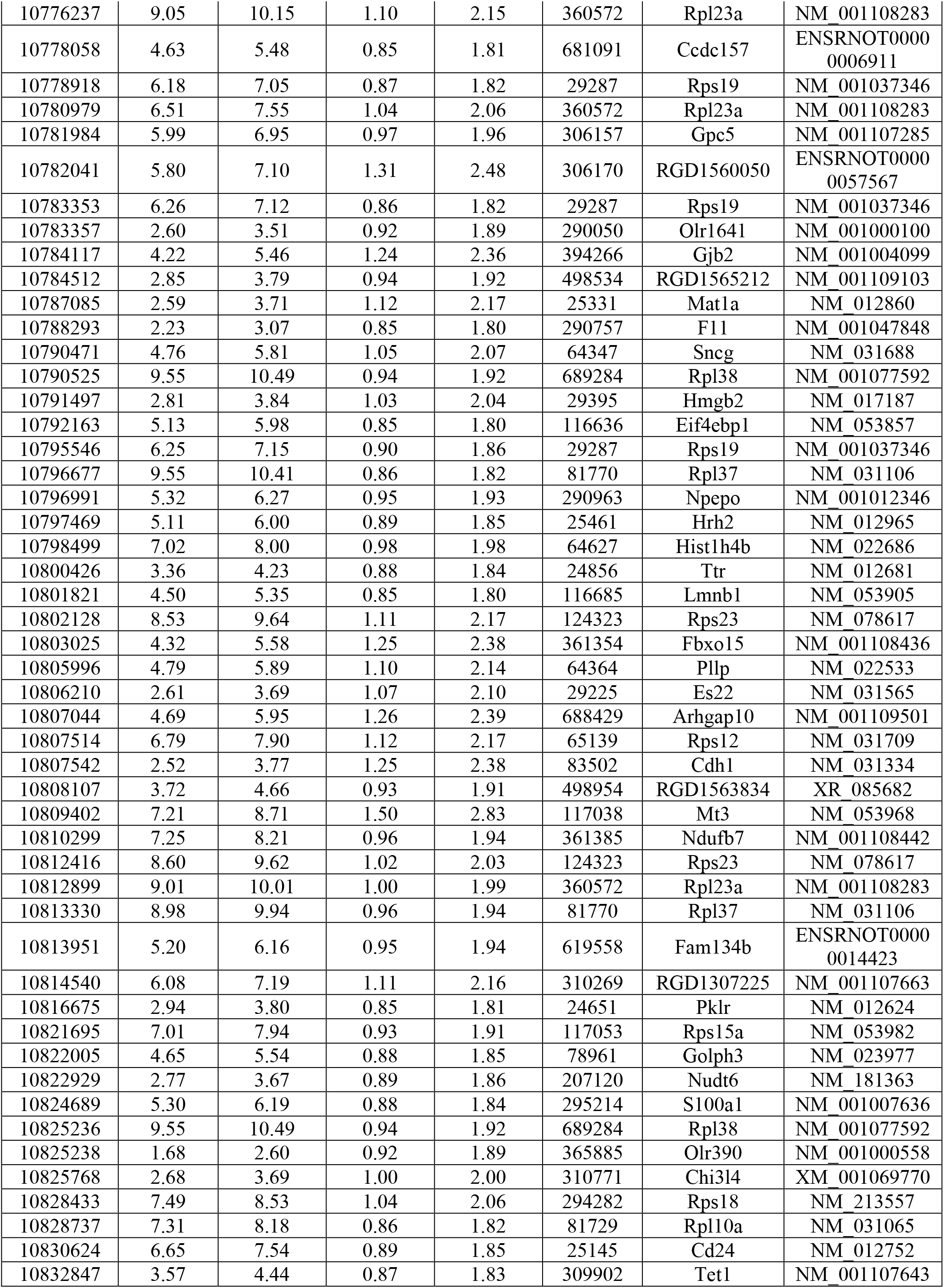
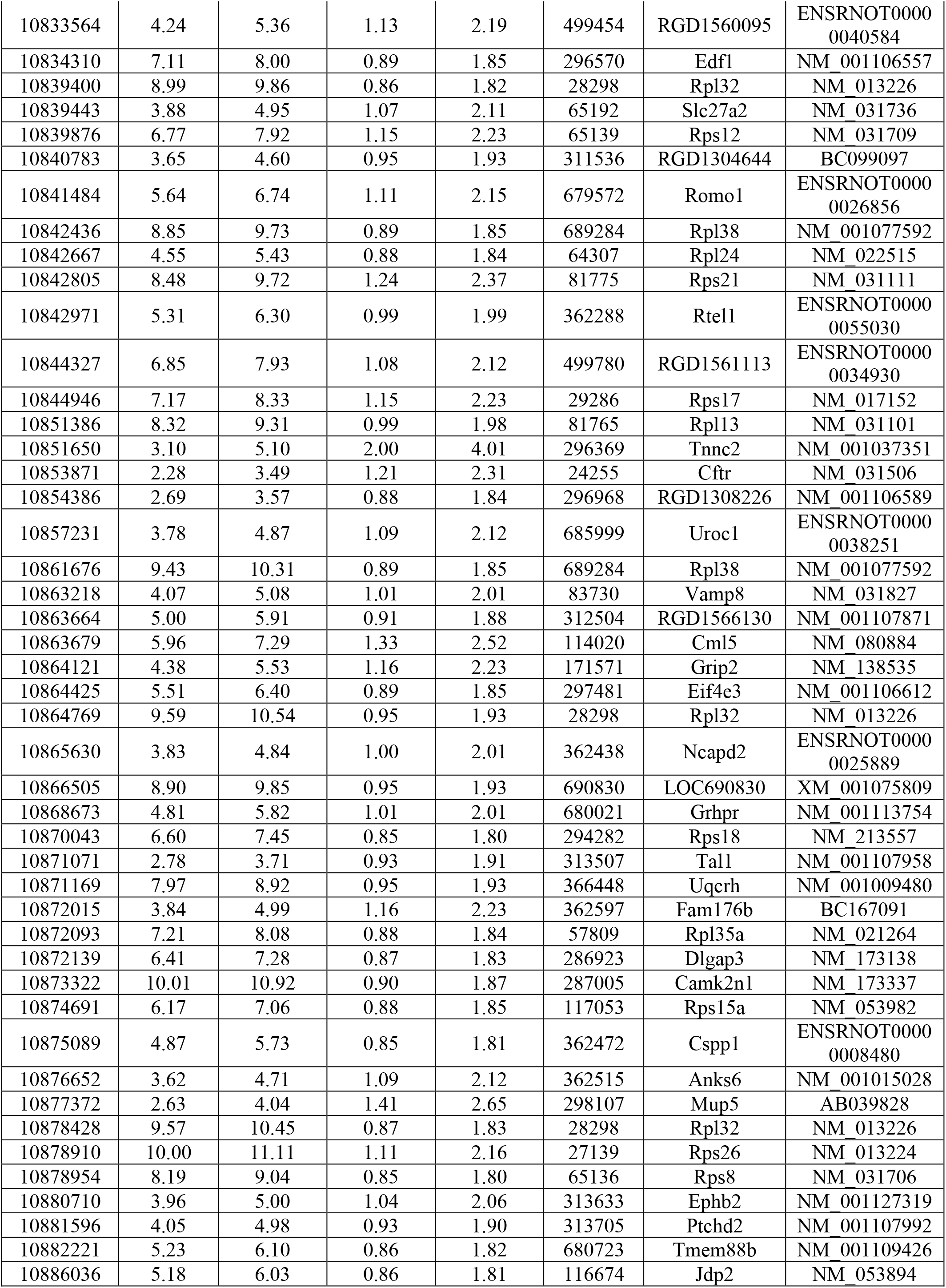
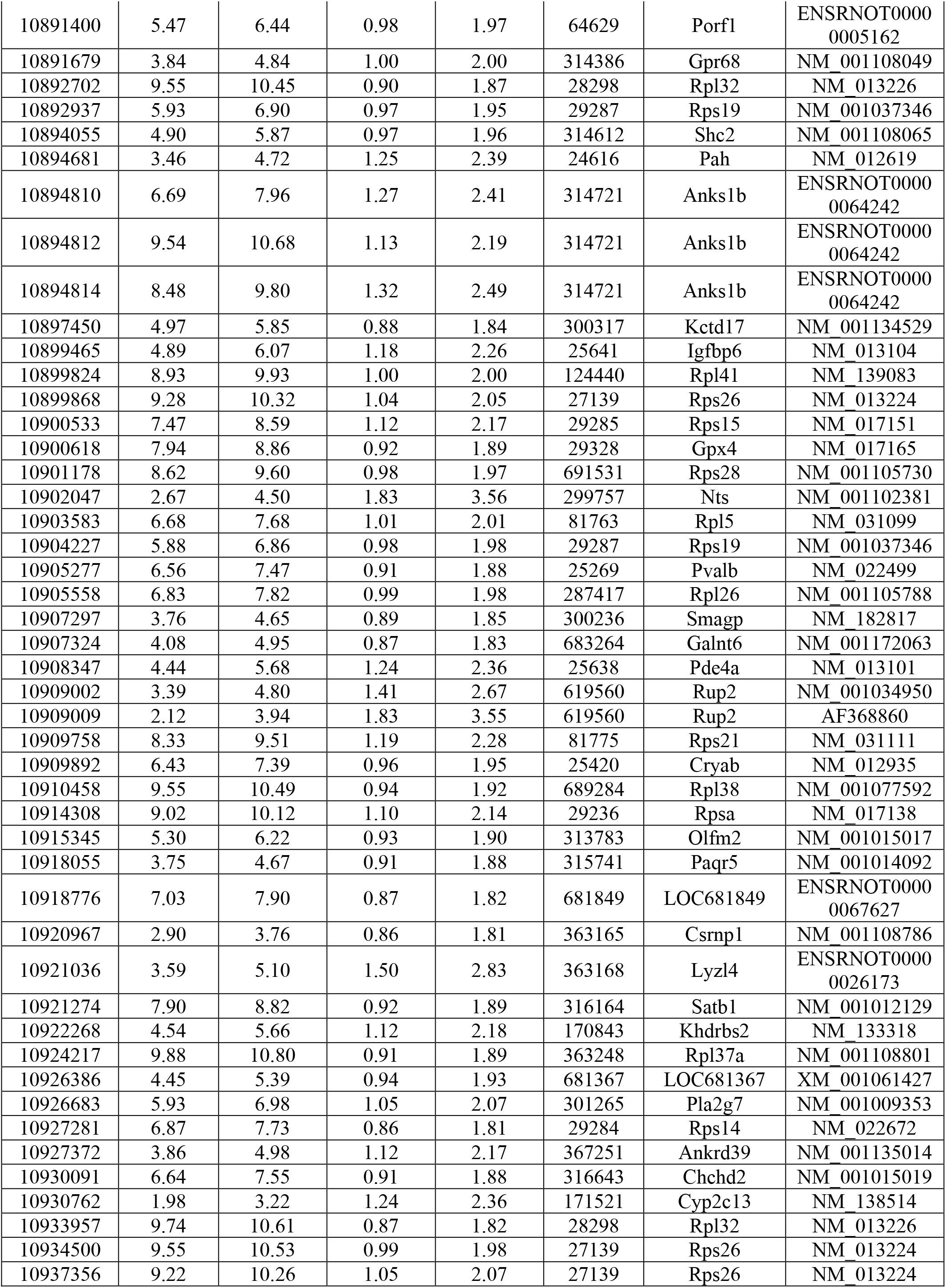
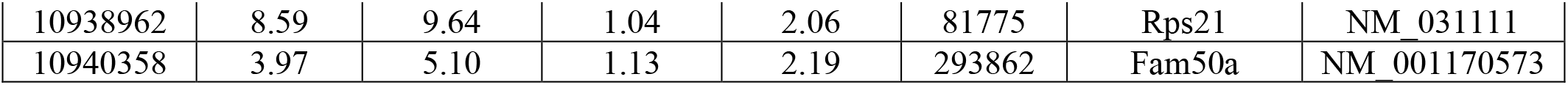
List of genes upregulated in brain.

**Supplementary table (ST-2B):**
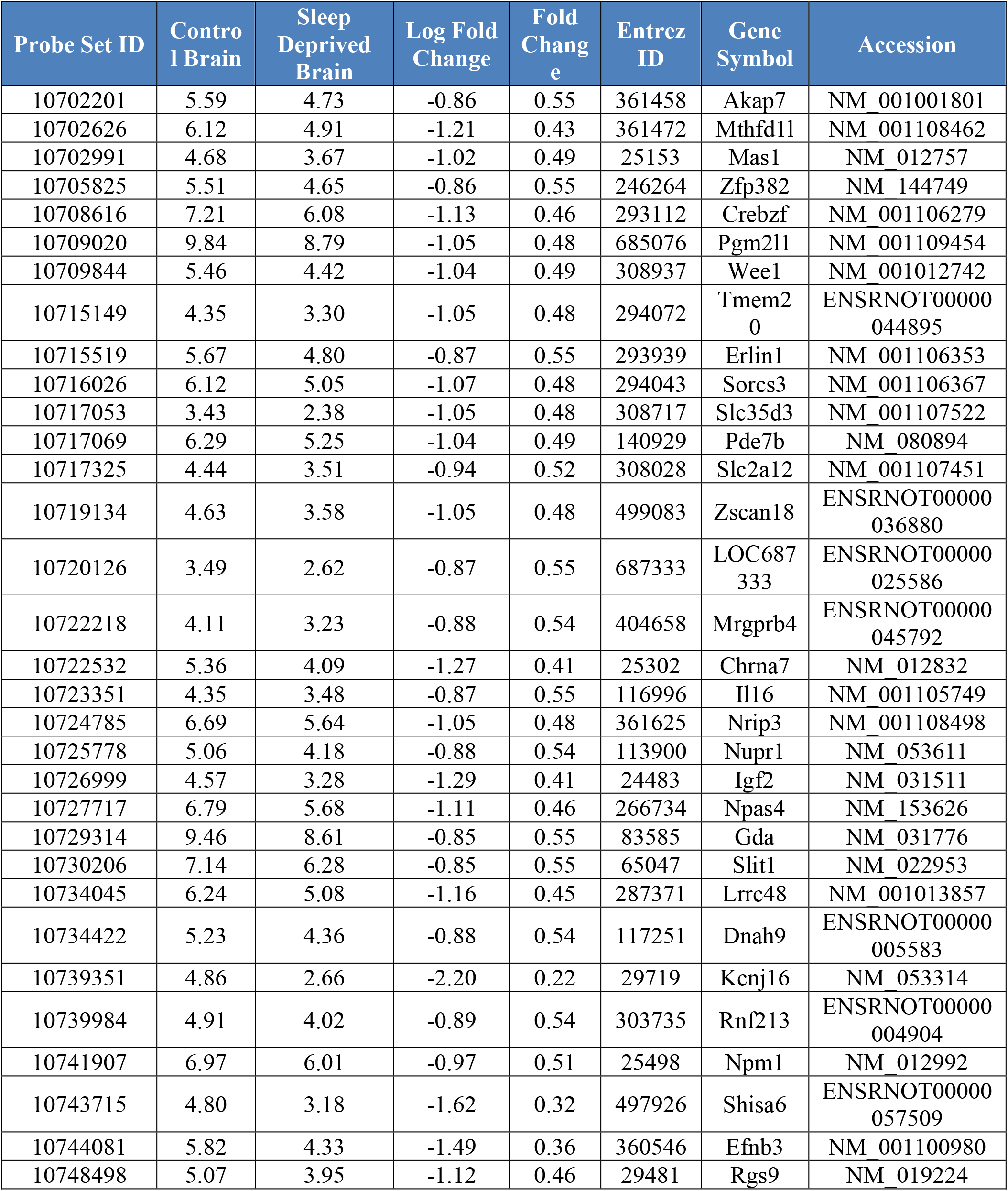
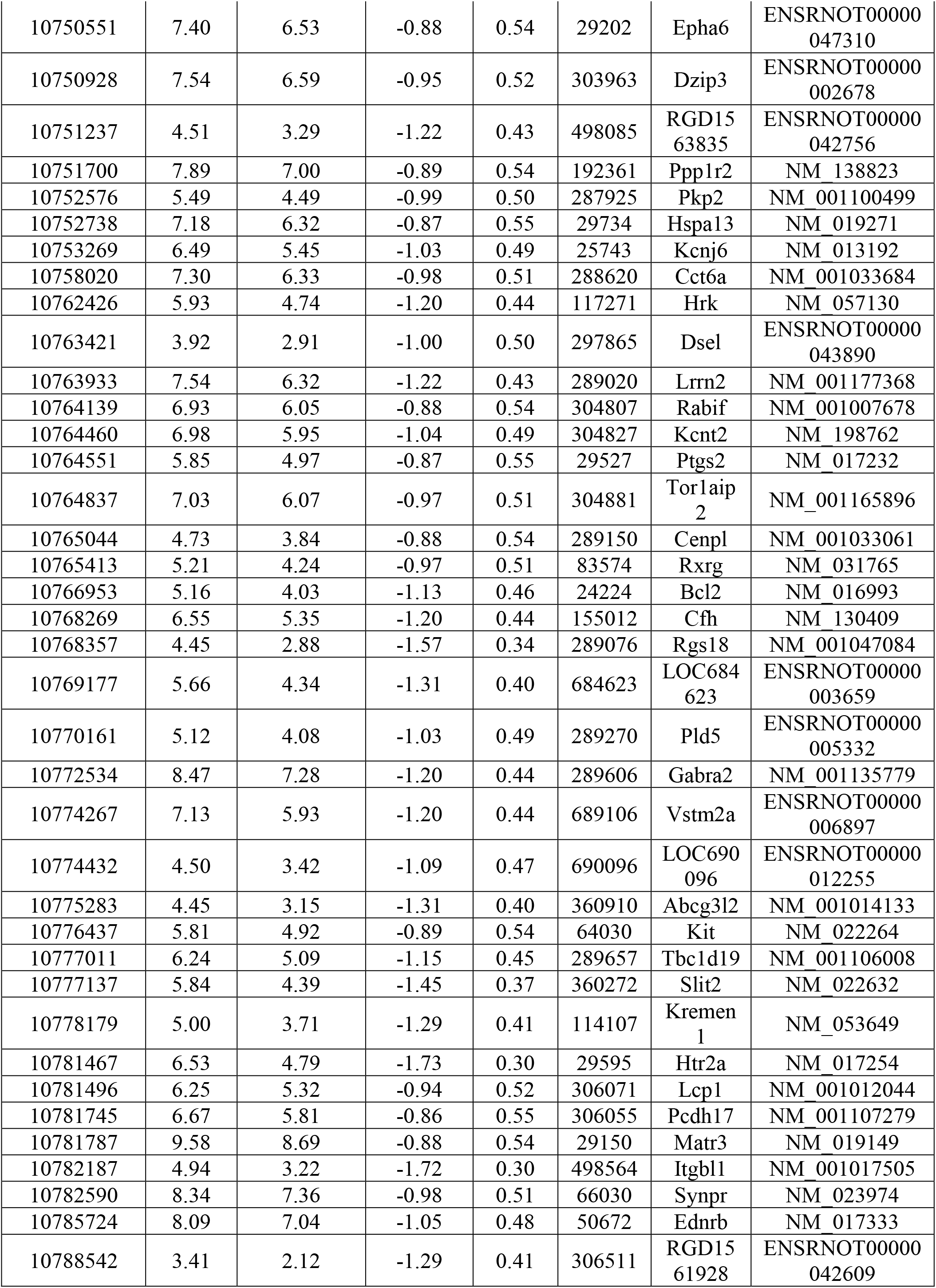
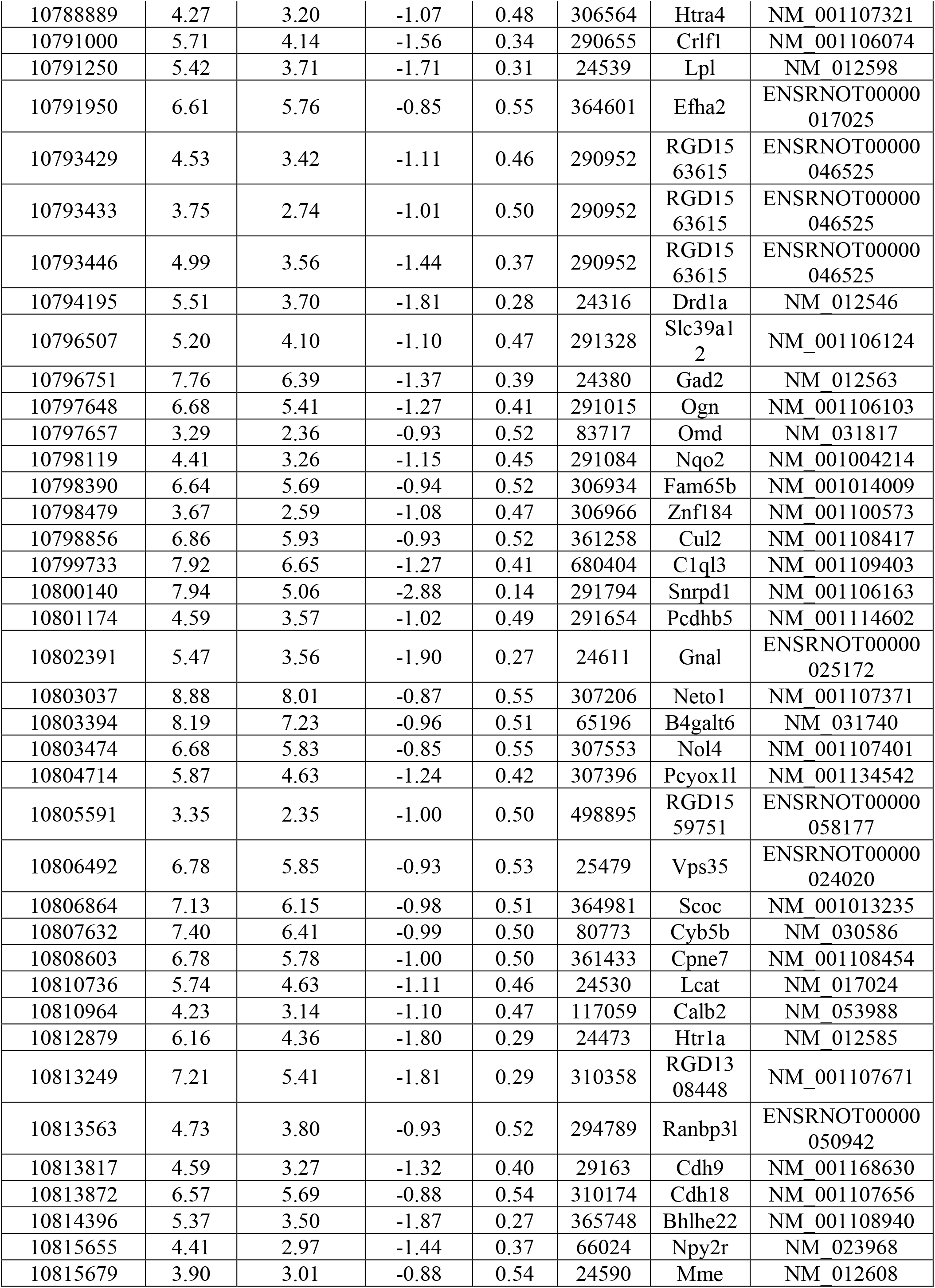
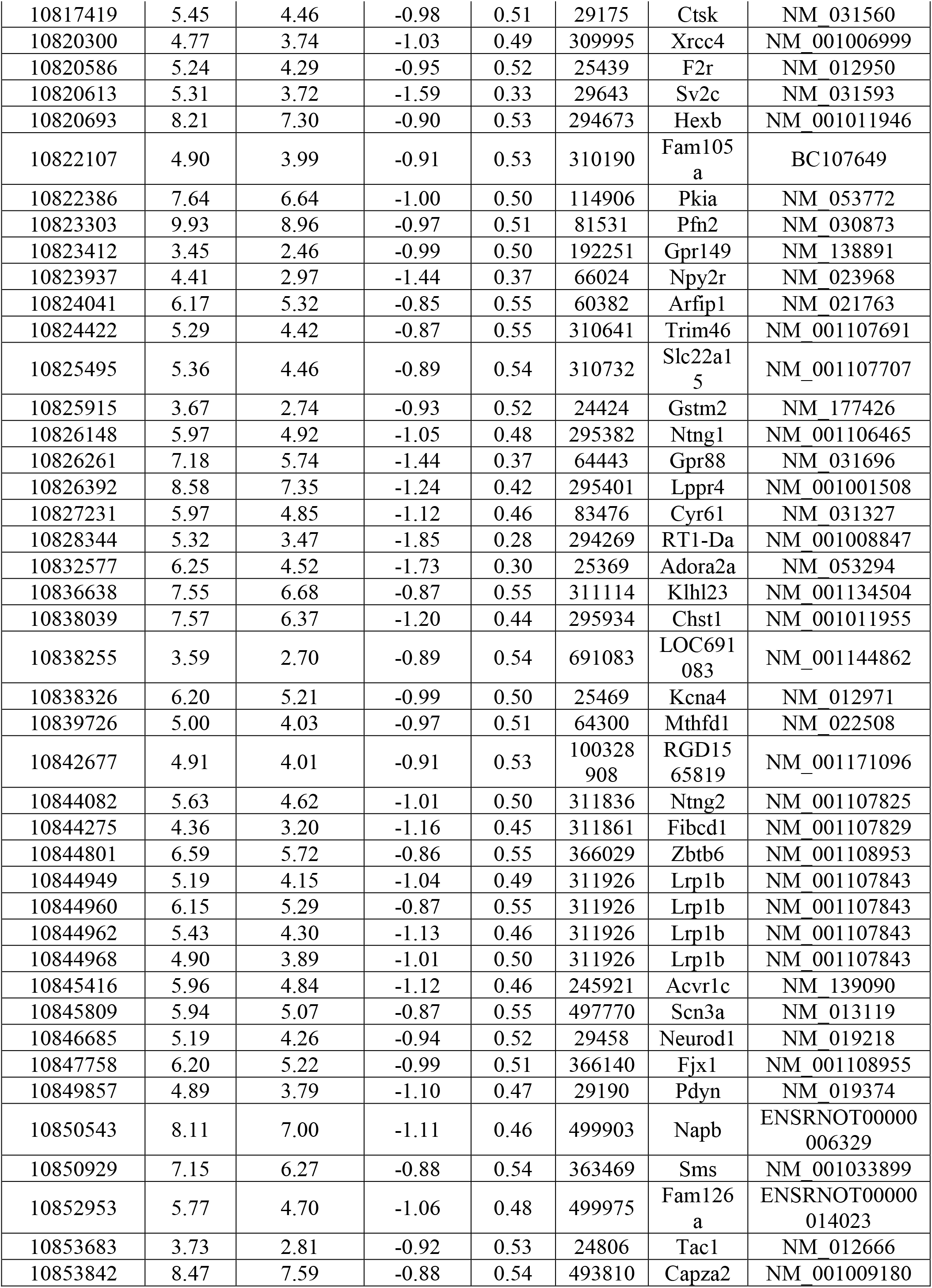
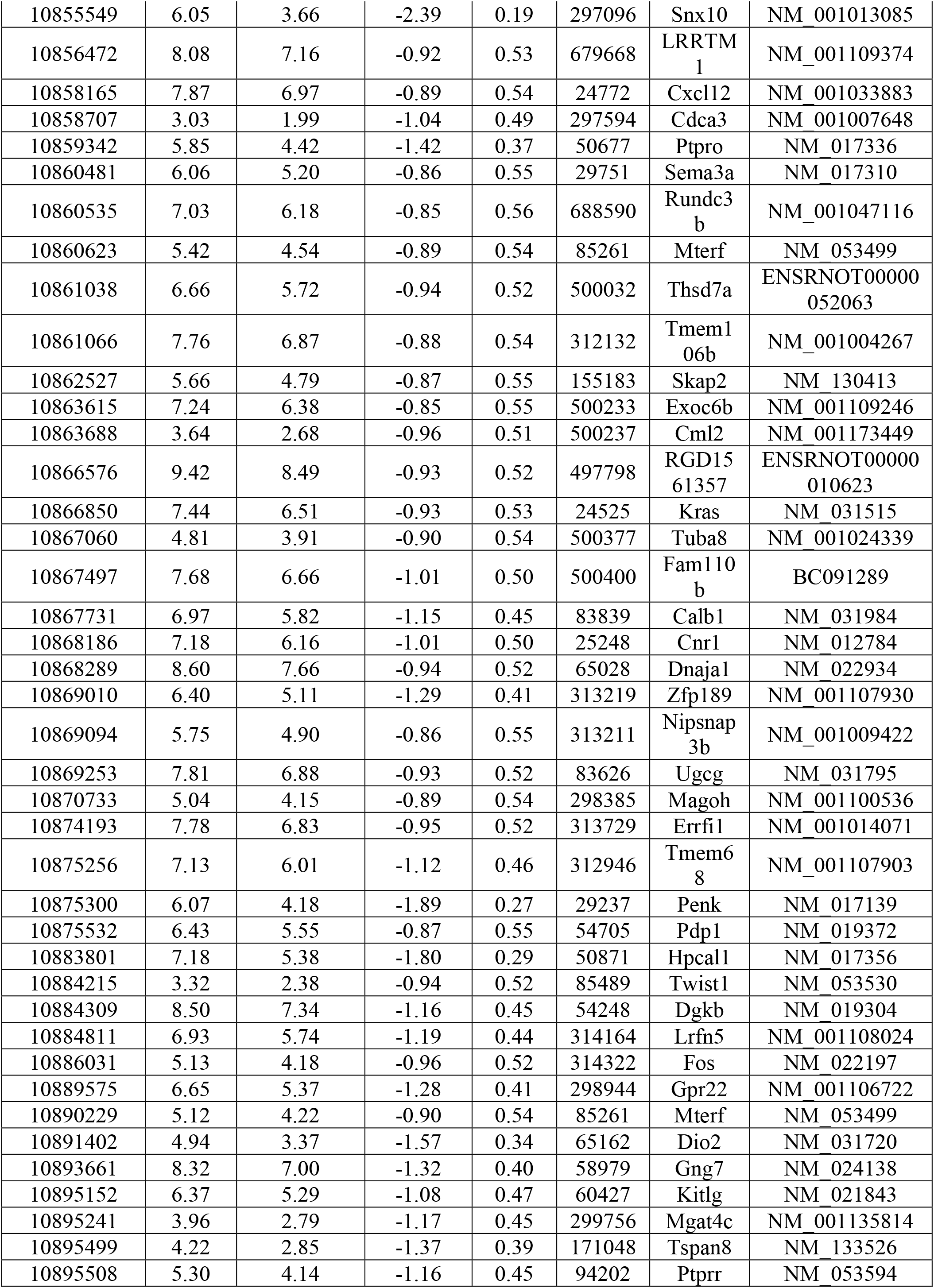
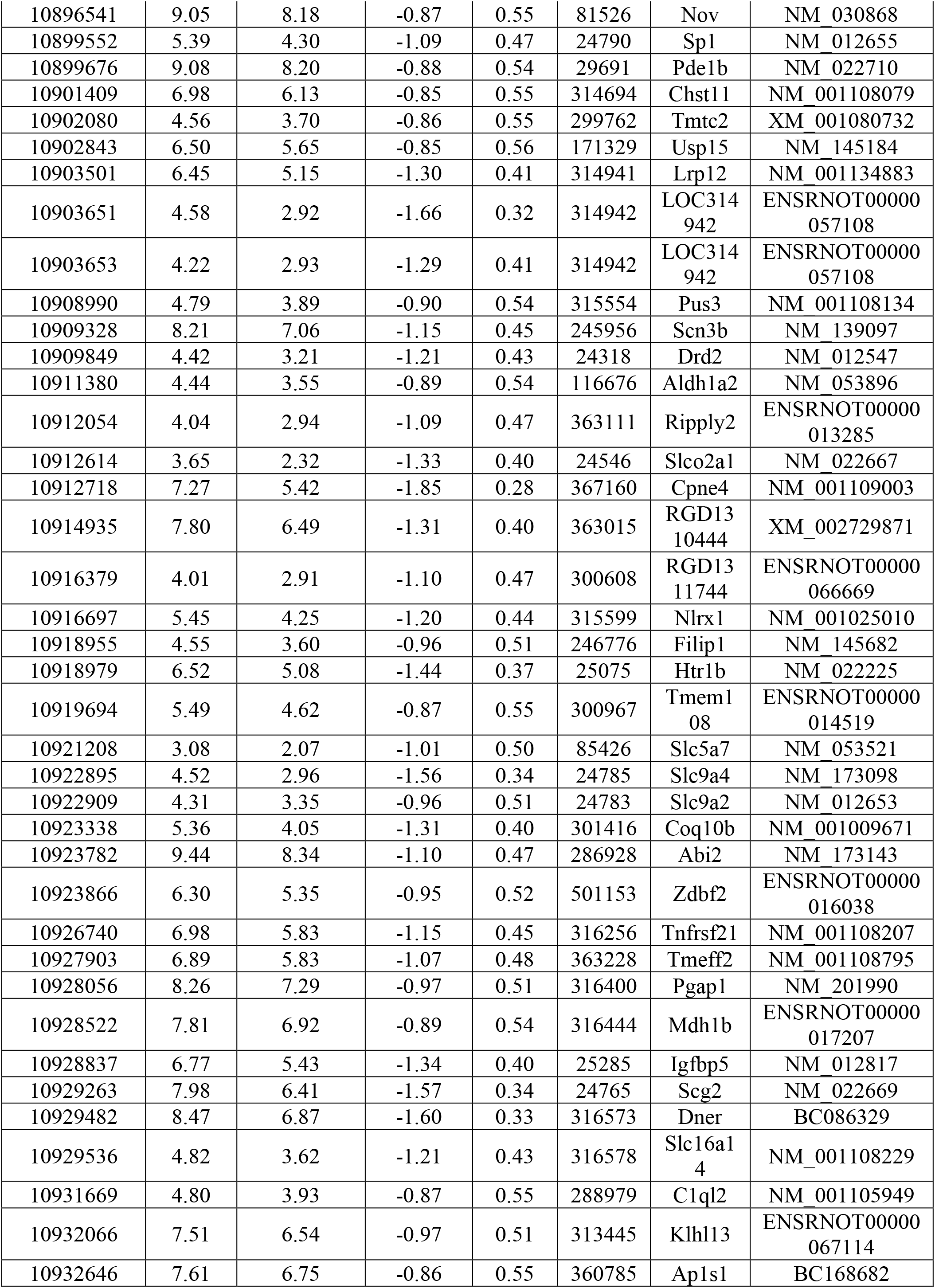
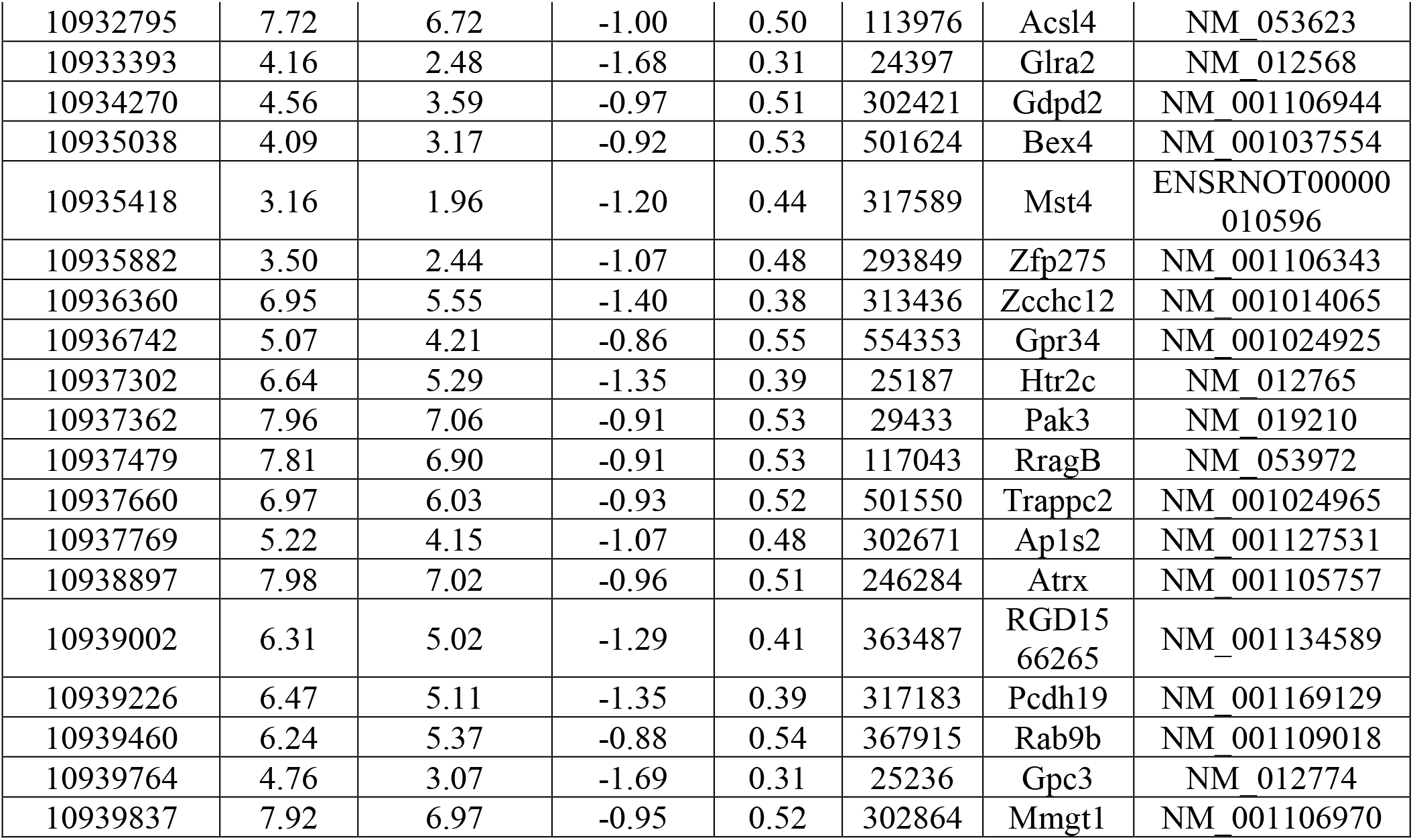
List of genes downregulated in brain.

**Supplementary table (ST-2C):**
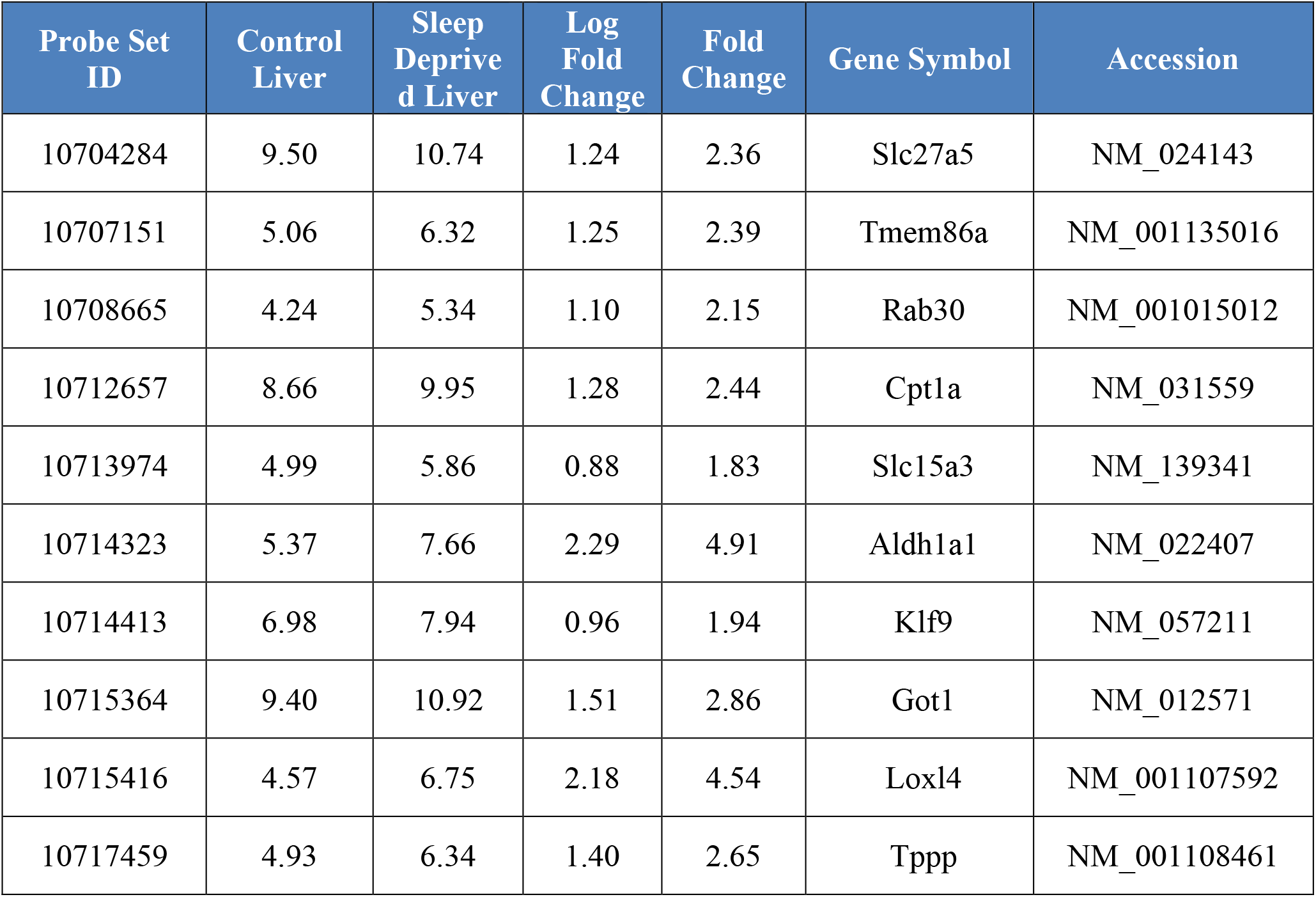
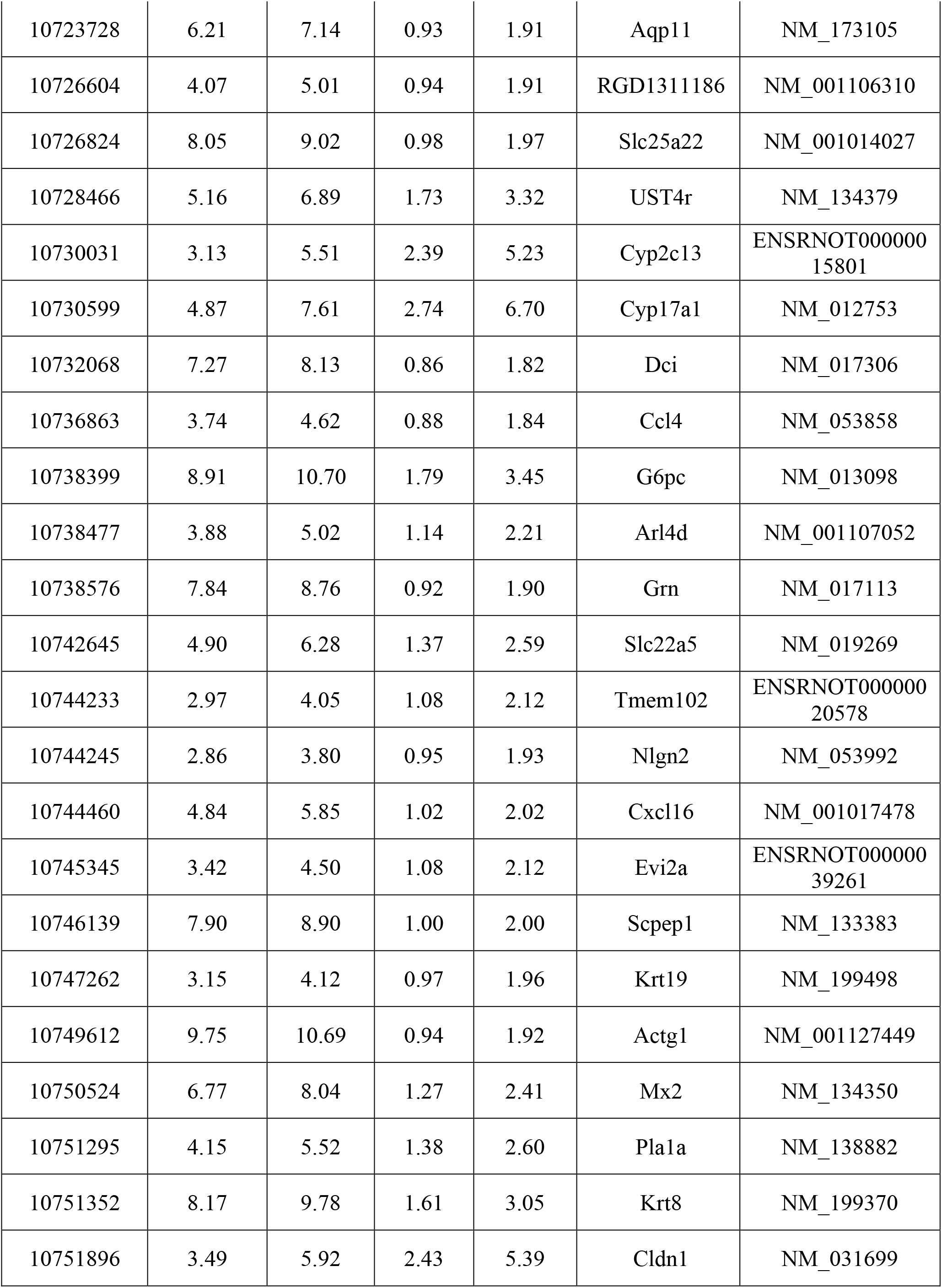
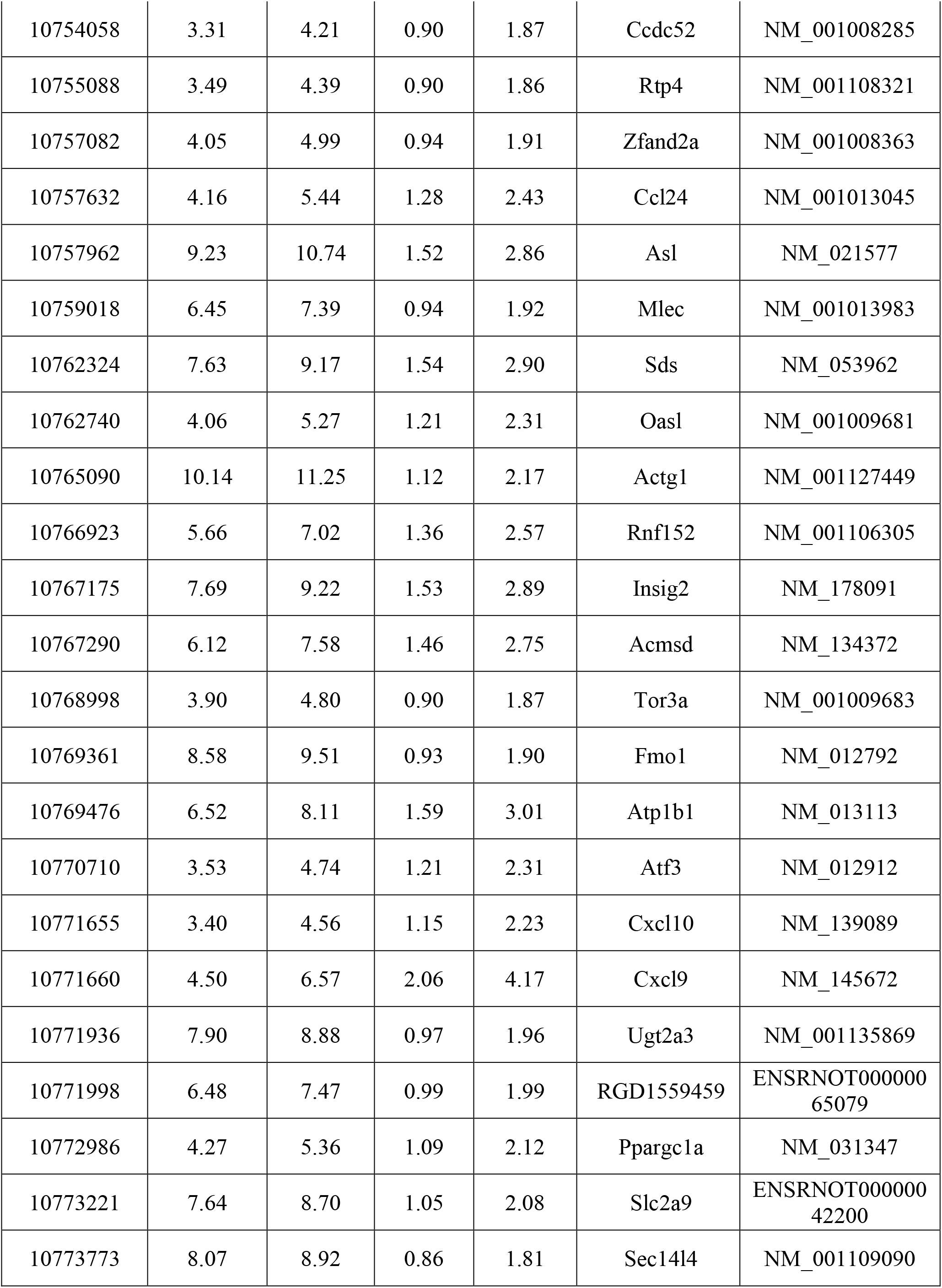
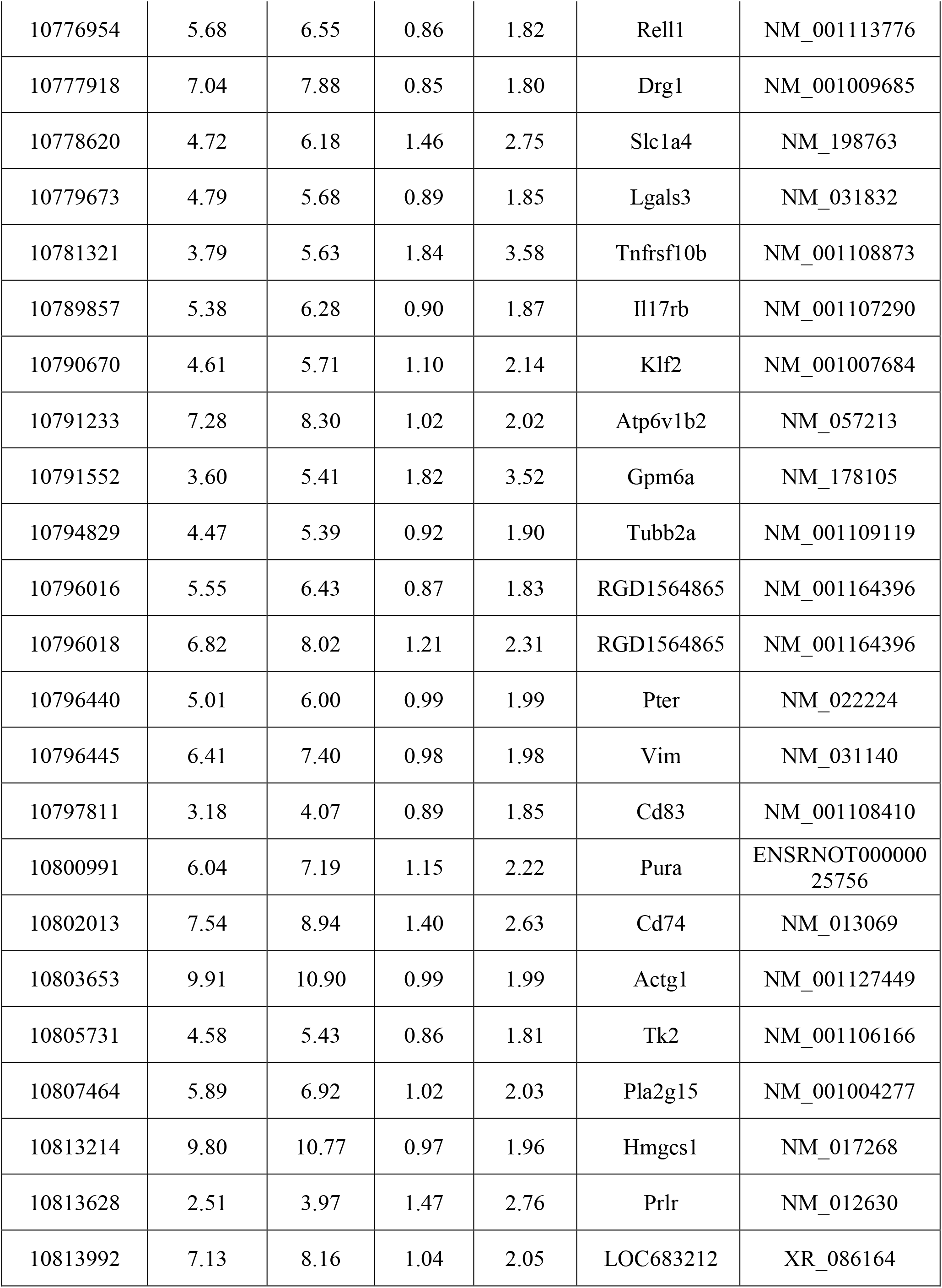
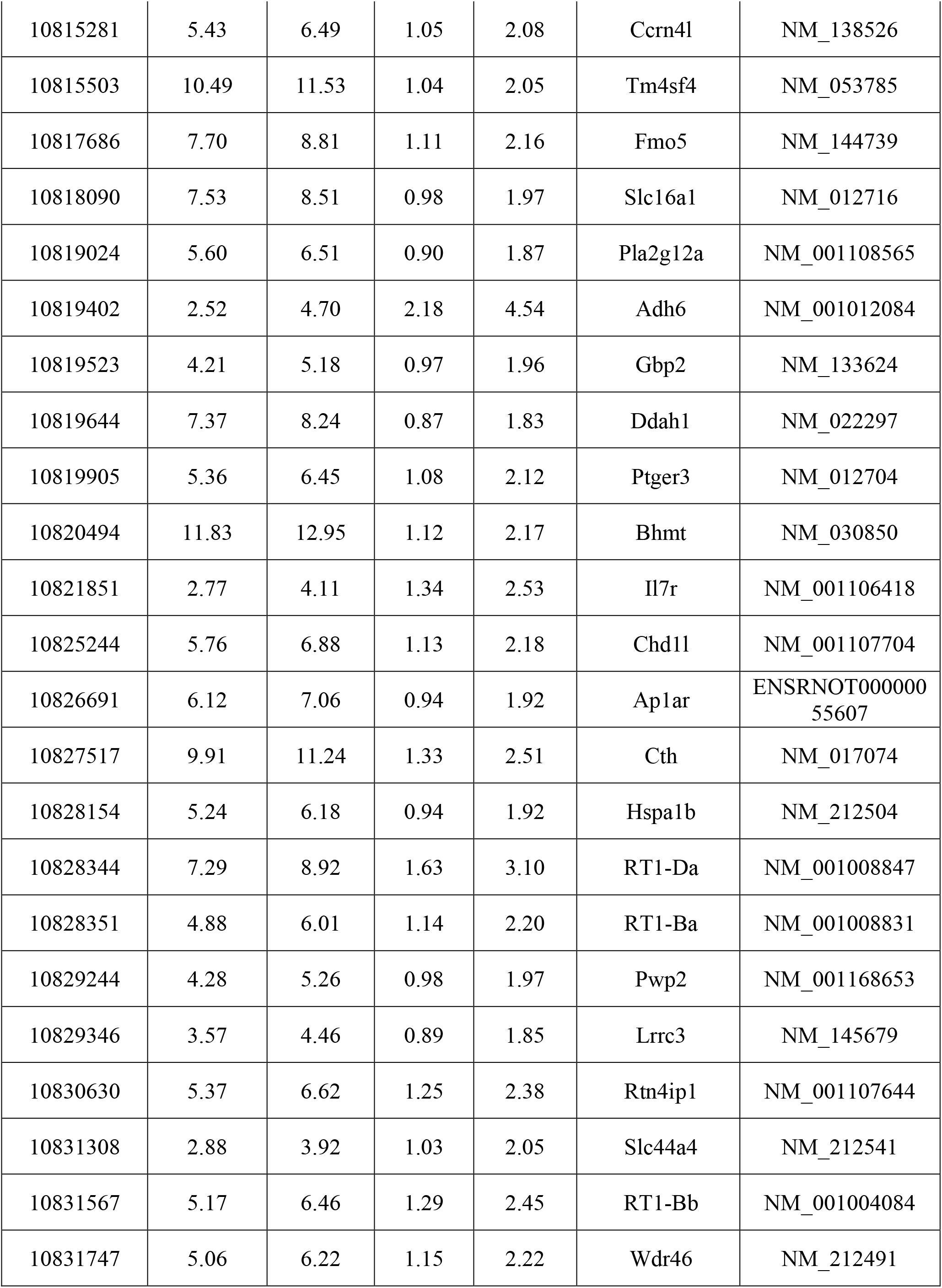
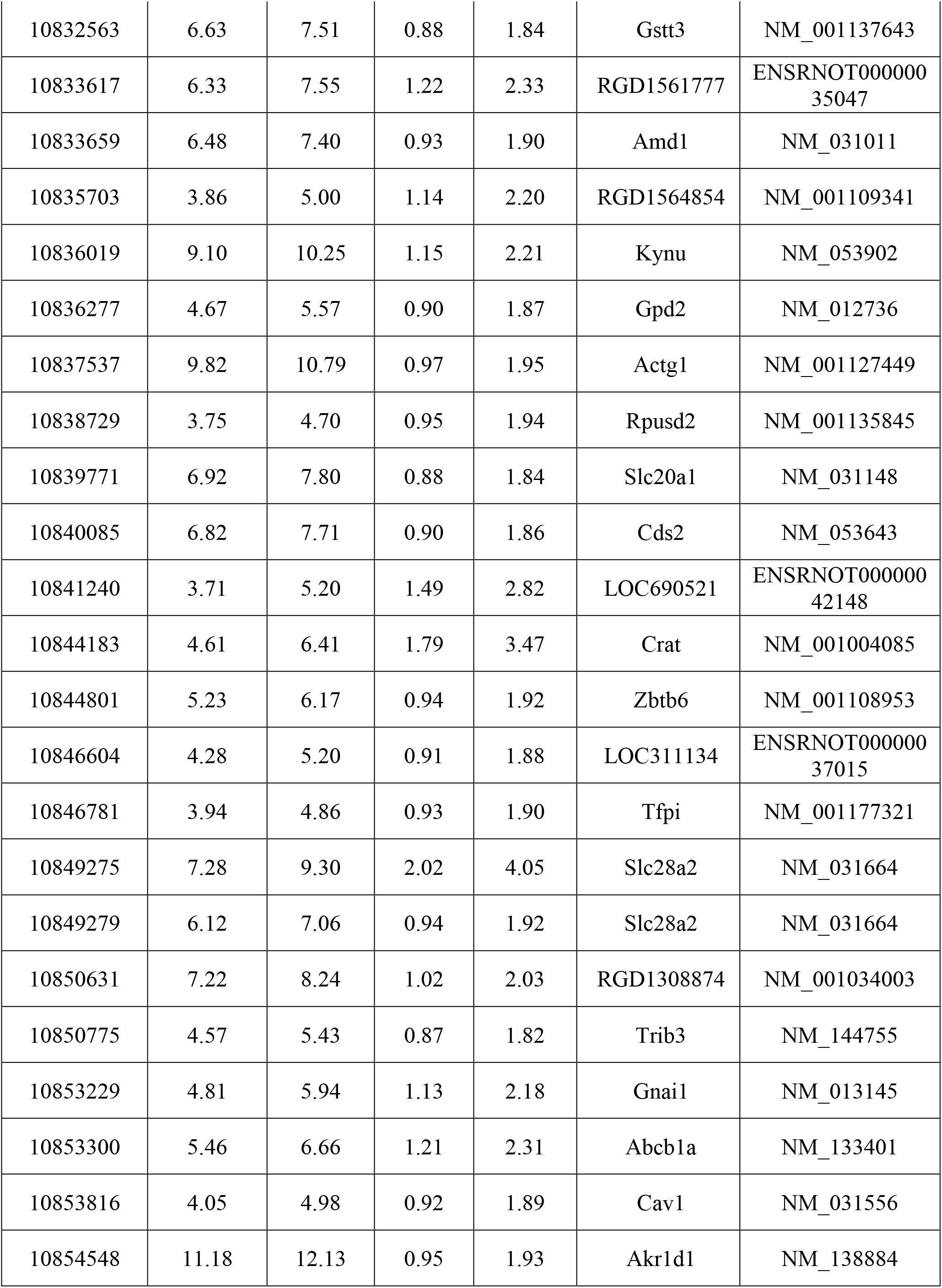
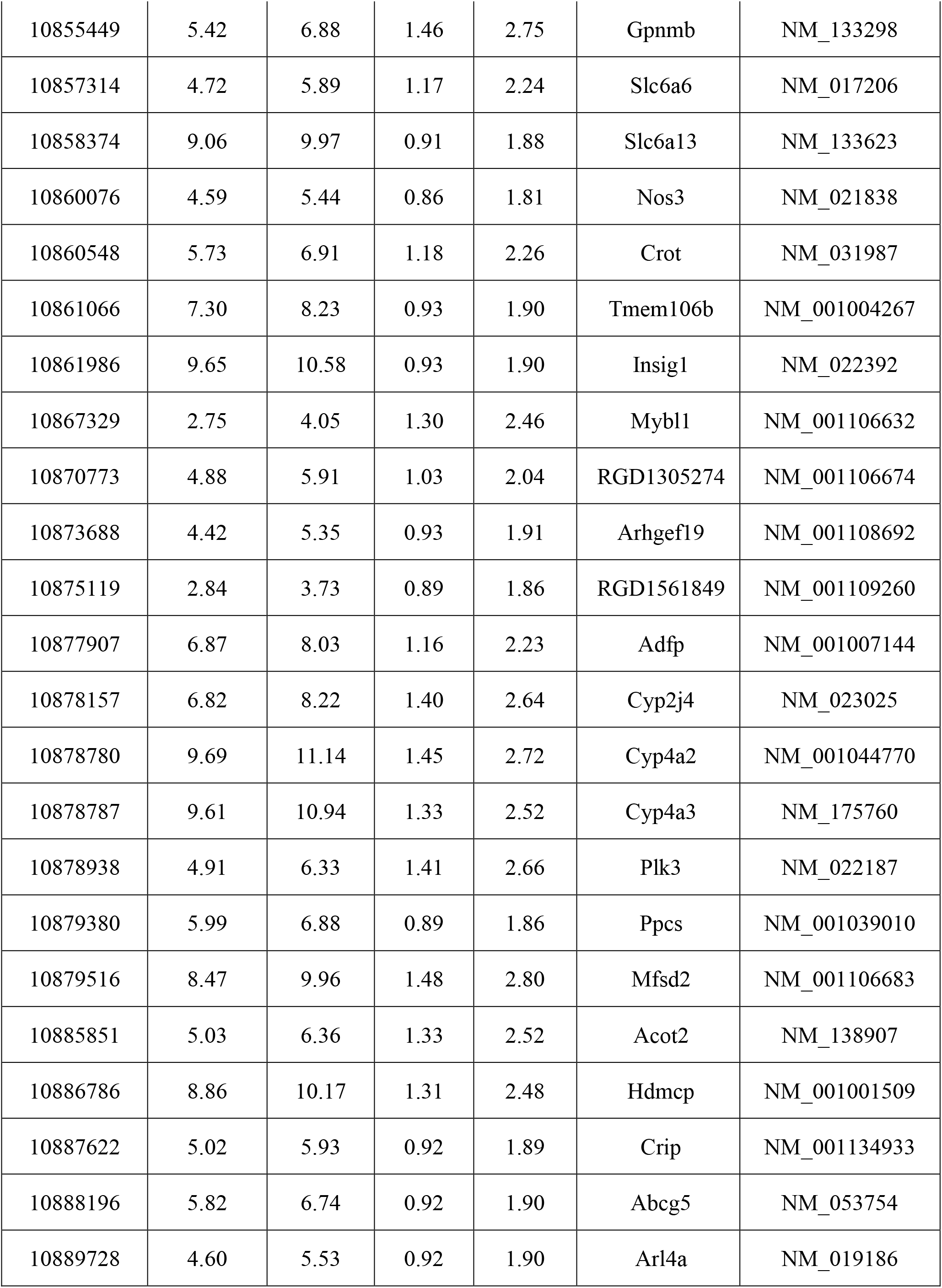
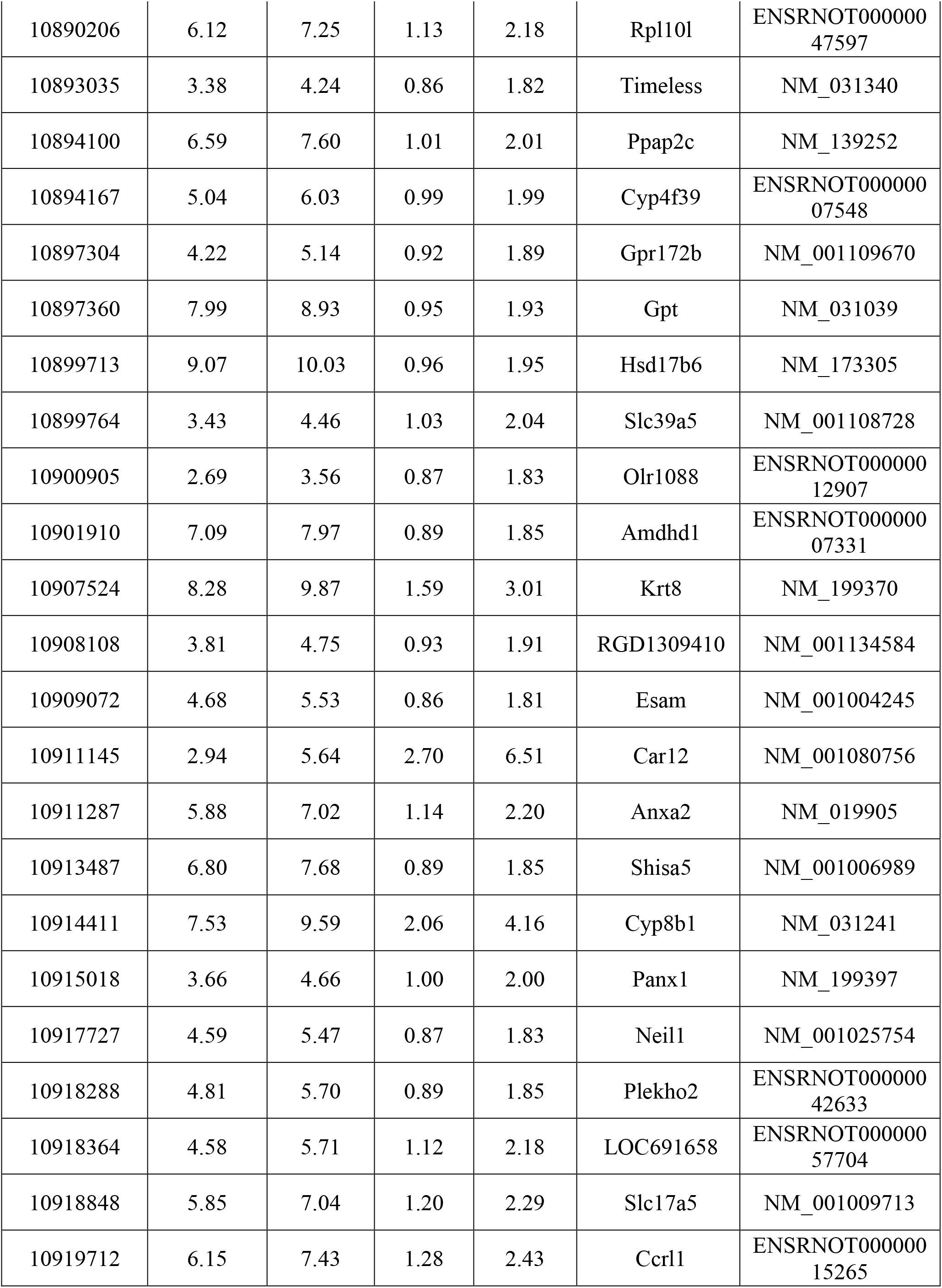
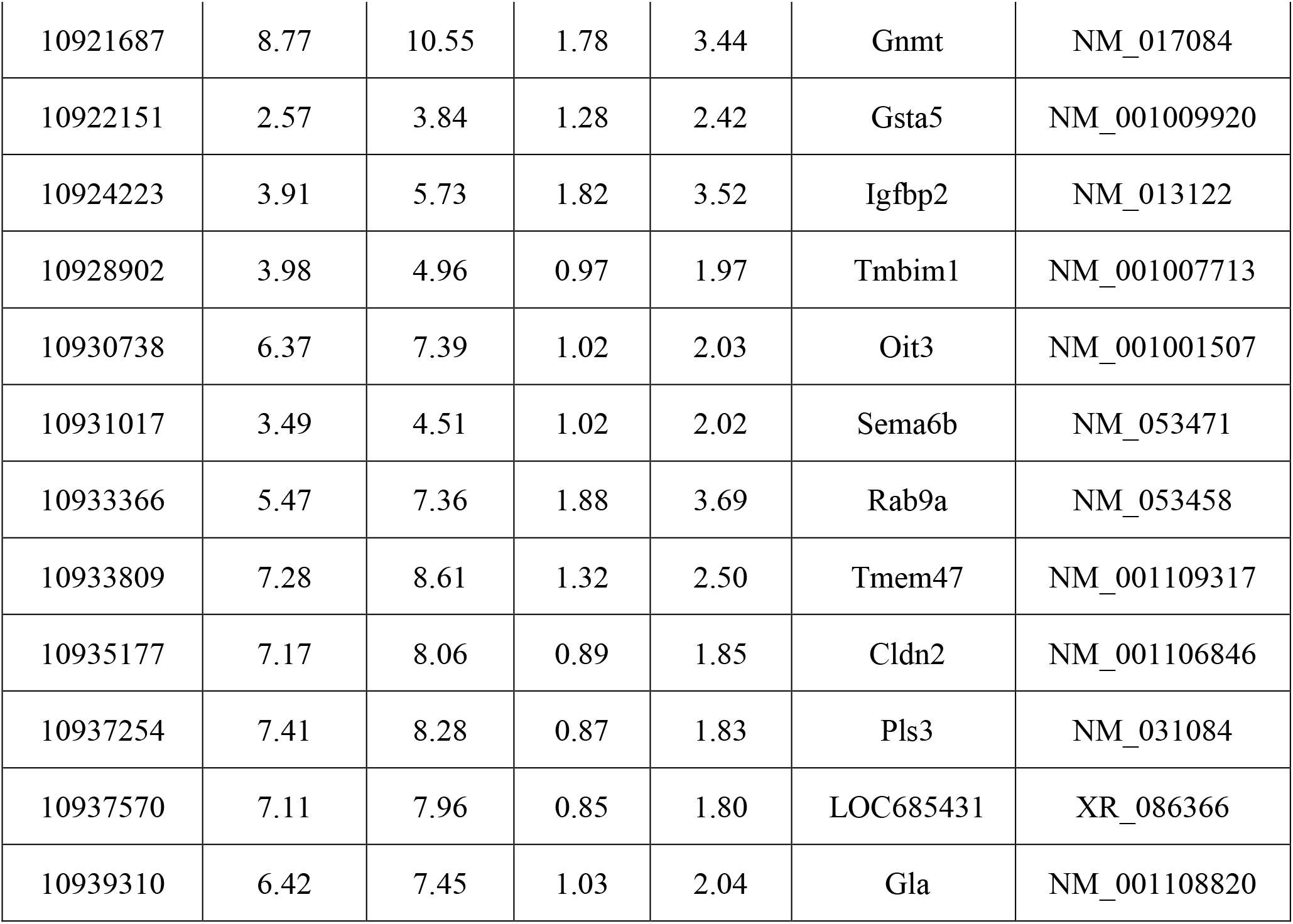
List of genes upregulated in liver.

**Supplementary table (ST-2D):**
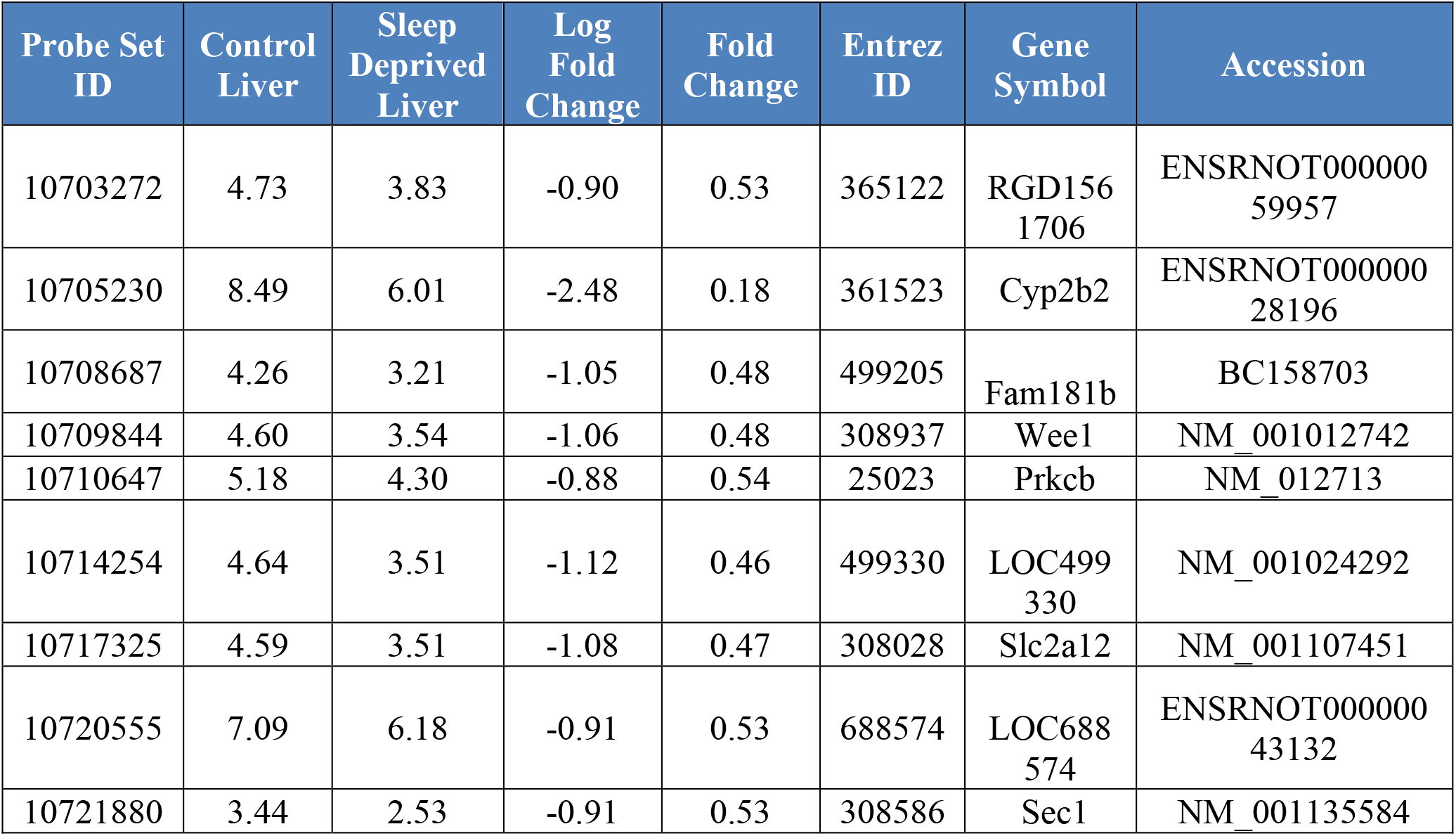
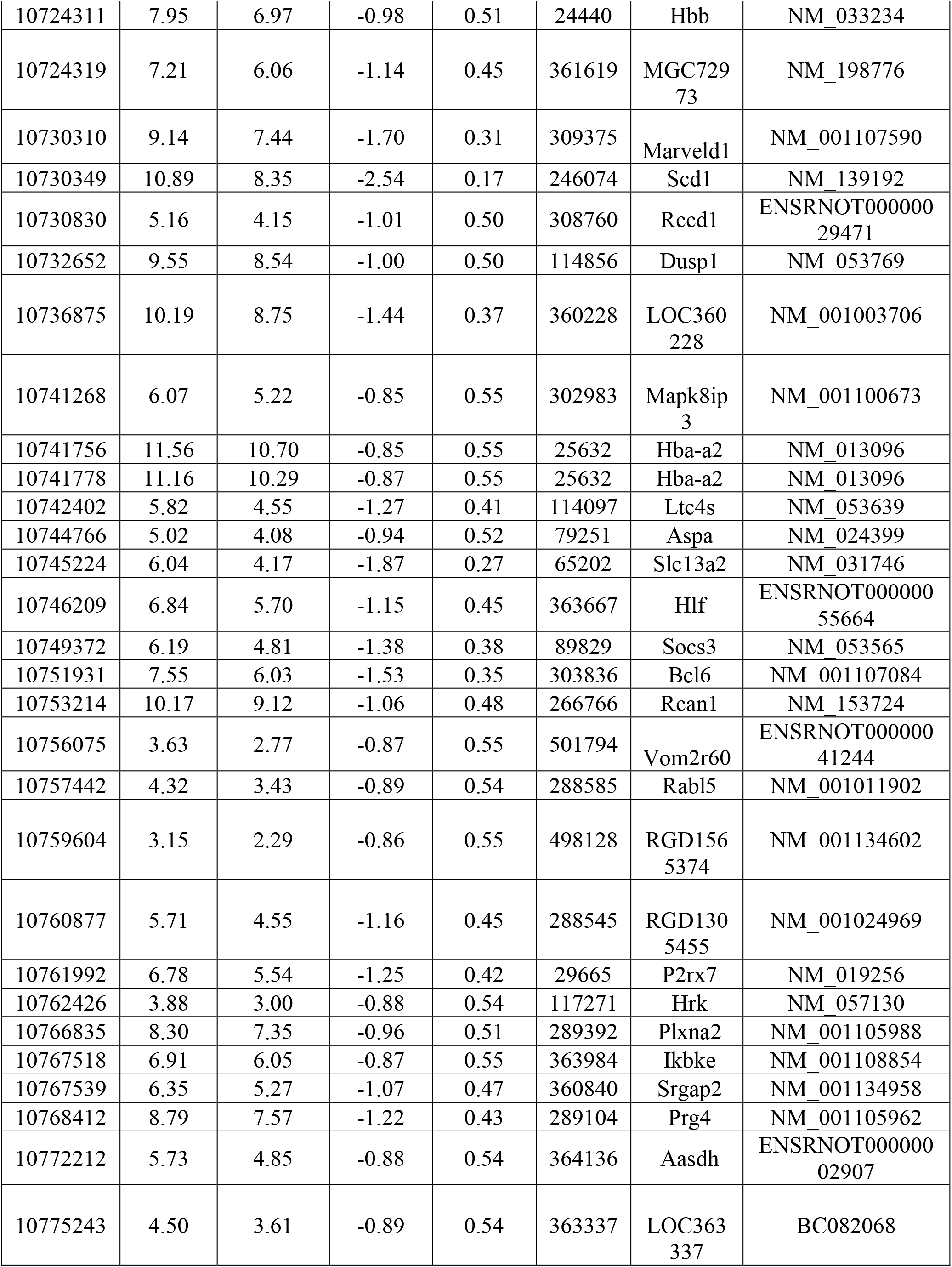
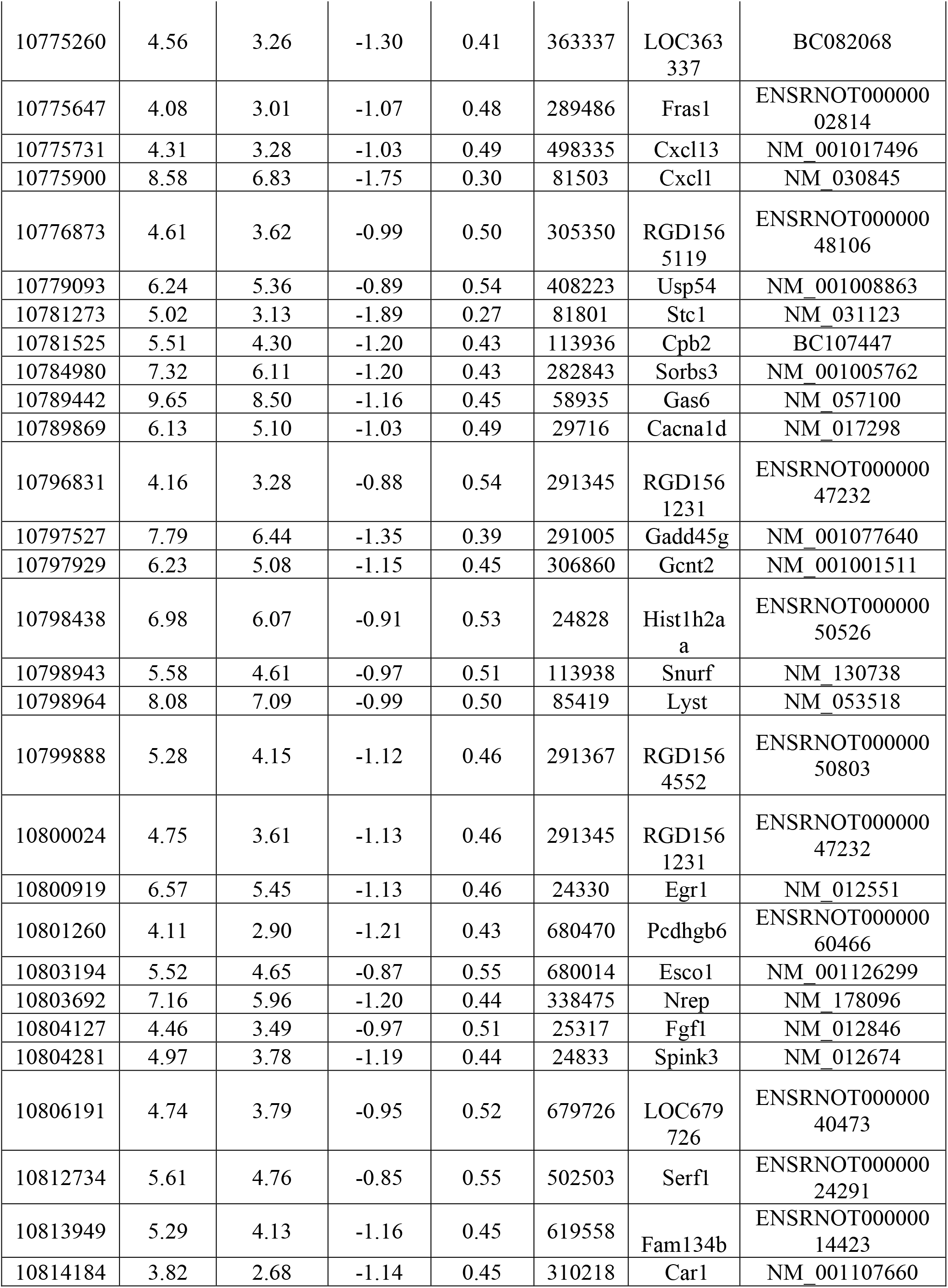
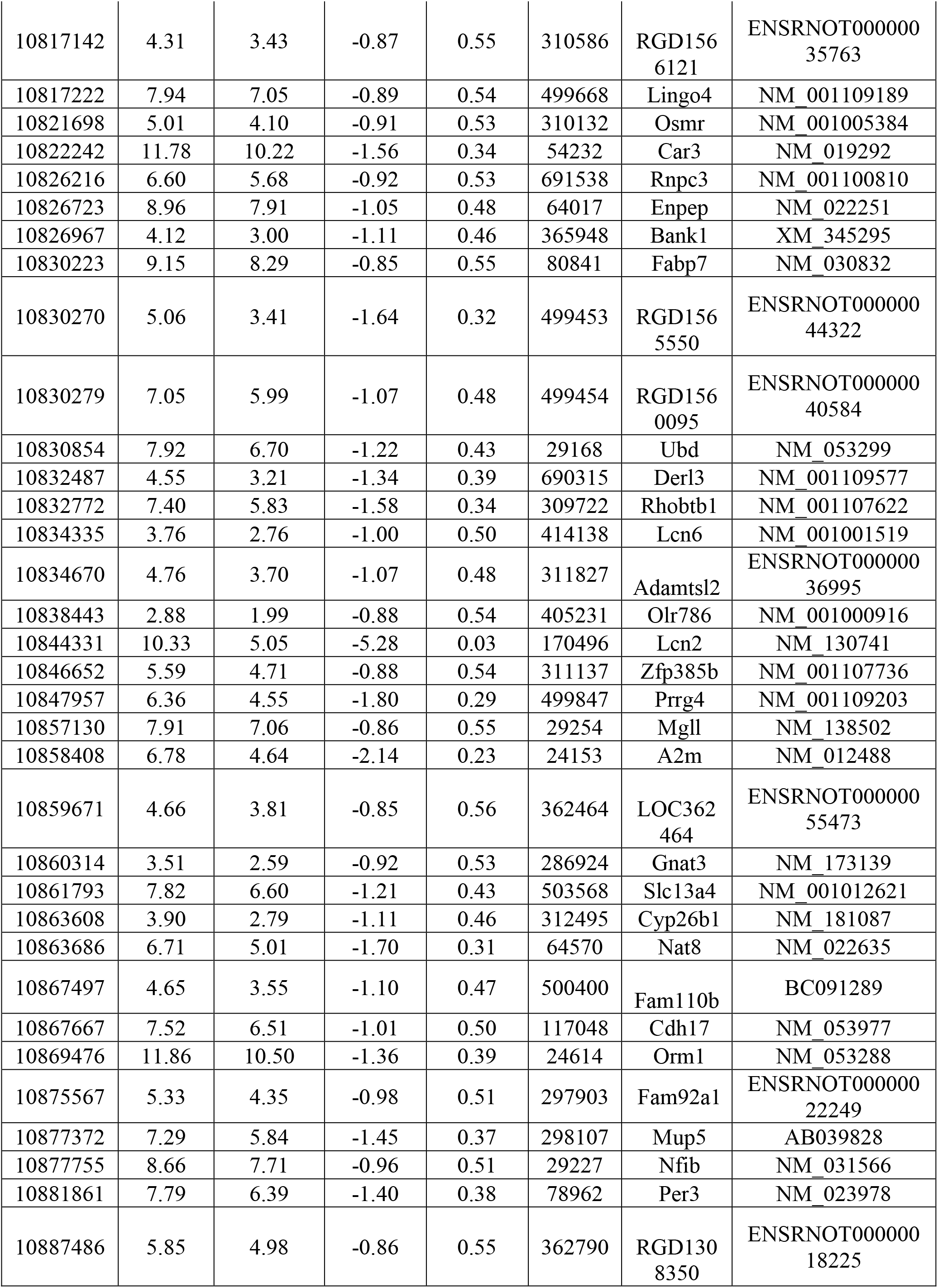
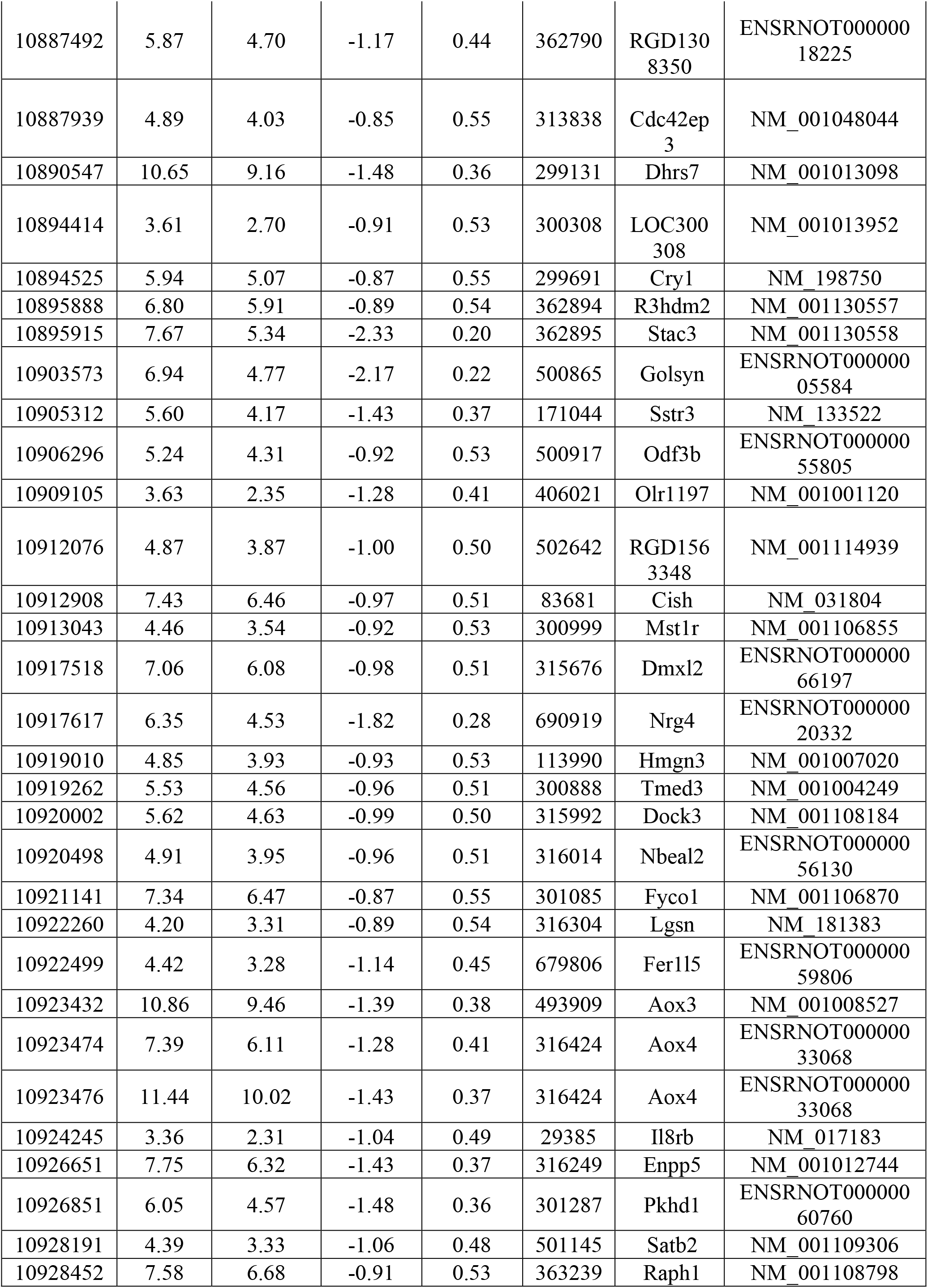
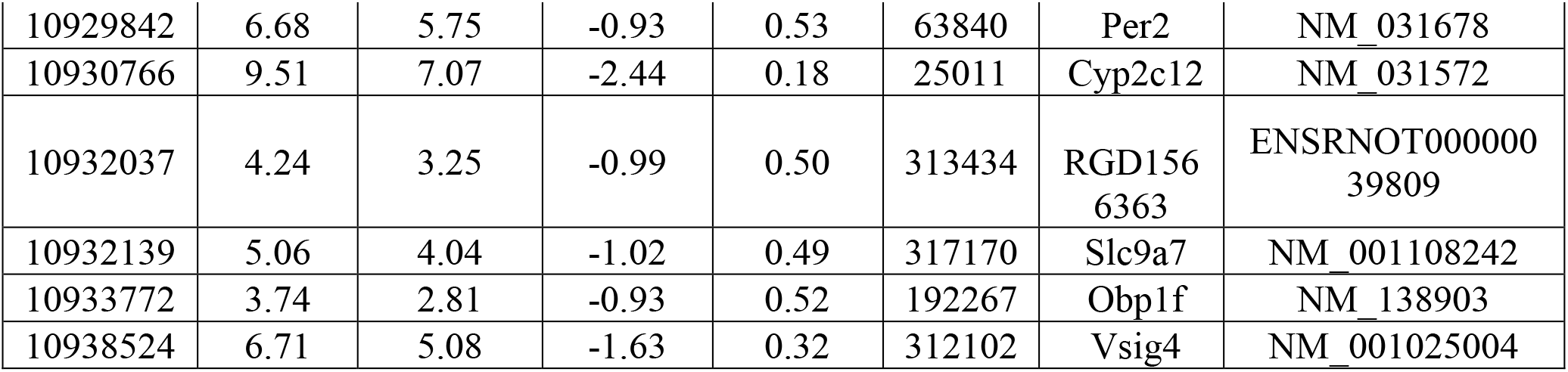
List of genes downregulated in liver.

**Supplementary table (ST3):**
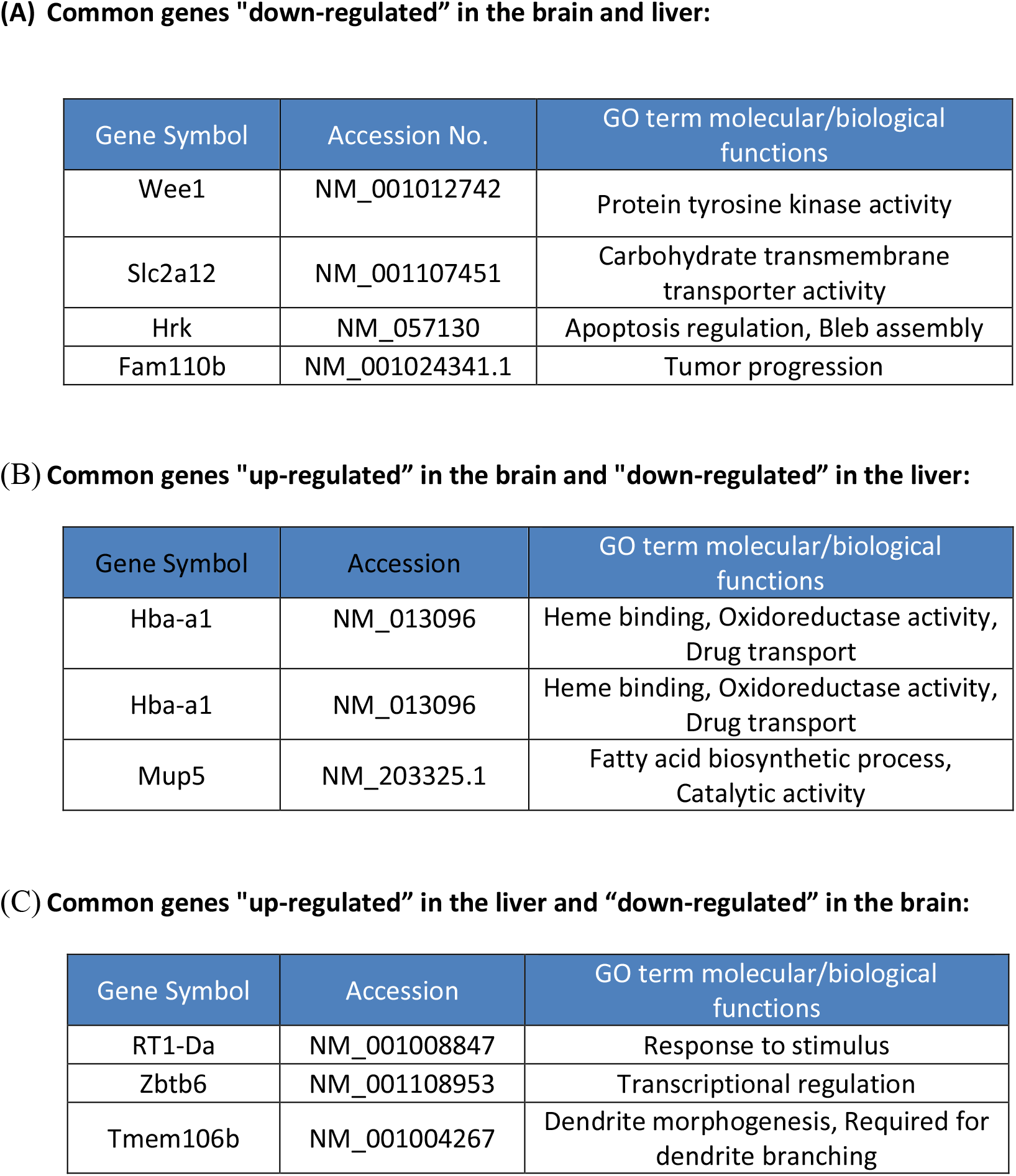
List of common genes affected in brain and liver.

**Supplementary Table 4 (ST-4):**
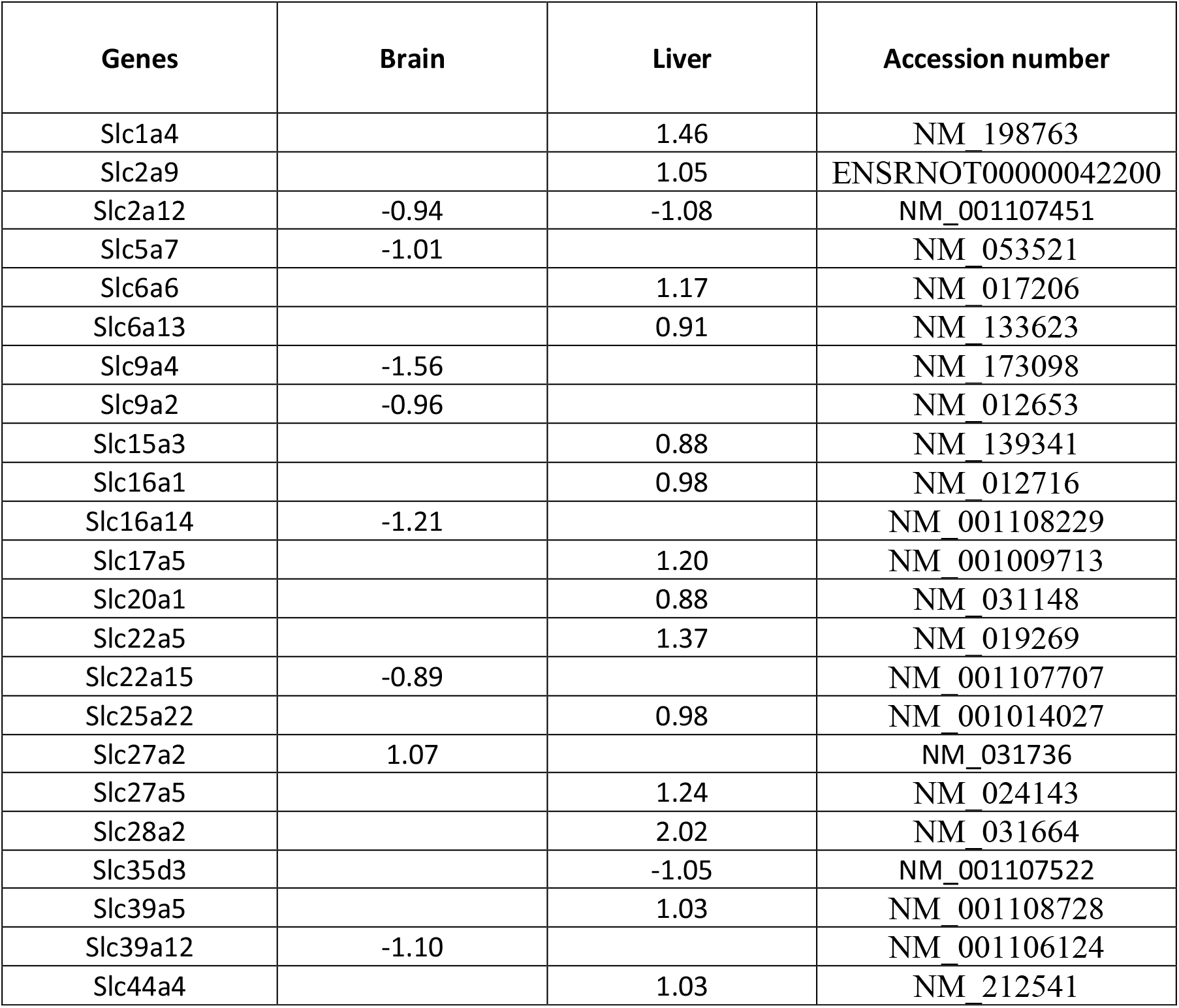
Log fold change of gene expression after REM sleep loss compared to control. Values in positive numbers shows upregulation while negative values show downregulation.

**Supplementary Table 5 (ST-5):** List of GO terms related to synaptic plasticity

| GO terms | Brain | RGD ID |
| --- | --- | --- |
| Positive regulation of synaptic plasticity | DR ($p=0.03$) | GO:0031915 |
| Positive regulation of long-term synaptic plasticity | UR ($p=0.002$) | GO:0048170 |
| Regulation of long-term neuronal synaptic plasticity | DR ($p=2.87E-005$) | GO:0048169 |
| Synaptic transmission | DR ($p= 0.001$) | |
| Synaptic transmission, dopaminergic | DR ($p= 2.64E-009$) | GO:0001963 |
| Regulation of synaptic transmission, dopaminergic | DR ($p= 0.0009$) | GO:0032225 |
| Negative regulation of synaptic transmission | DR ($p= 0.004$) | GO:0050805 |
| Negative regulation of synaptic transmission, glutamatergic | DR ($p=7.77E-009$) | GO:0051967 |
| Positive regulation of synaptic transmission, glutamatergic | DR ($p=9.93E-006$) | GO:0051968 |
| Synaptic transmission, cholinergic | DR ($p= 1.33E-006$) | GO:0007271 |
| Regulation of synaptic transmission, GABAergic | DR ($p=2.47E-006$) | GO:0032228 |
| Positive regulation of synaptic transmission, GABAergic | DR ($p=0.0002$) | GO:0032230 |
| Positive regulation of synaptic transmission, glutamatergic | DR ($p=9.93E-006$) | GO:0051968 |
| Negative regulation of synaptic transmission, glutamatergic | DR ($p=7.77E-009$) | GO:0051967 |
| Long term synaptic depression | DR ($p=0.006$) | GO:0060292 |
| Long-term synaptic potentiation | DR ($p=0.009$) | GO:0060291 |
| Neuromuscular synaptic transmission | DR ($p=0.03$) | GO:0007274 |
| Negative regulation of synaptic transmission, dopaminergic | DR ($p= 0.04$) | GO:0032227 |
| Establishment of synaptic specificity at neuromuscular junction | DR ($p=0.04$) | GO:0007529 |

**Supplementary Table 6 (ST-6):** List of GO terms related to immune system

| GO terms | Brain | Liver | RGD ID |
| --- | --- | --- | --- |
| Immune response | | UR ($p=0.0002$) | GO:0006955 |
| Immunoglobulin mediated immune response | | UR ($p=0.009$) | GO:0016064 |
| Negative regulation of type 2 immune response | | DR ($p=0.001$) | GO:0002829 |
| Regulation of immune system process | | DR ($p=0.01$) | GO:0002682 |
| Type 2 immune response | | DR ($p=0.01$) | GO:0042092 |
| Positive regulation of T cell mediated immune response to tumor cell | UR ($p=0.0006$) | | GO:0002842 |
| Immune system development | DR ($p=0.001$) | | GO:0002520 |
| T cell activation involved in immune response | DR ($p=0.002$) | | GO:0002286 |

## Notes

### Competing Interest Statement

The authors have declared no competing interest.

### Summary of Updates

We revised the article after peer-reviews comments and changed the figures.

